# From promiscuity to specialization: The origin of cholinergic and monoaminergic signaling is pre-bilaterian

**DOI:** 10.64898/2026.10.13.758597

**Authors:** Alexandros A. Pittis, Luis Alfonso Yañez-Guerra, Fabian Ruperti, Alison G. Cole, Milena Marinković, Daniel Thiel, Jaime Huerta-Cepas, Jacob M. Musser, Ulrich Technau, Gáspár Jékely, Detlev Arendt

## Abstract

The emergence of chemical neurotransmission was key to the evolution of nervous systems, yet when the different transmitter systems arose remains unclear. Biogenic amines (here, acetylcholine and the monoamines) have been reported in non-bilaterian animals and even outside Metazoa, but their physiological role has only recently begun to be resolved, and phylogenetic surveys of the bilaterian machinery have restricted dedicated monoaminergic signaling to Bilateria. Combining targeted metabolomics, molecular phylogenies, structural modeling and binding-affinity prediction, single-cell expression analysis and immunochemistry, we show that cholinergic and monoaminergic signaling predate Bilateria. We detect acetylcholine, dopamine, histamine, adrenaline, tyramine, 2-phenylethylamine and tryptamine in a sponge and two cnidarians, and identify broad-specificity enzymes and transporters able to produce and load them. These genes are co-expressed in both neuronal and non-neuronal cell types, indicating that amine signaling was never confined to the nervous system. Two *Hydra vulgaris* muscarinic receptor orthologs retain a fully conserved orthosteric site and are predicted to bind acetylcholine as strongly as their bilaterian counterparts, and muscle-type nicotinic receptor subunits are co-expressed with Rapsyn-related genes in cnidarian muscle cells, placing cholinergic neuromuscular signaling in the cnidarian–bilaterian ancestor. Specialization came later in Bilateria, where gene duplications produced substrate-specific enzymes (ChAT, TH, TPH), dedicated vesicular transporters (VAChT, VMAT) and ligand-selective receptors. Biogenic amine signaling is thus not a bilaterian innovation but an ancient mode of cellular communication that evolved from generalist to specialist machinery.

## Introduction

Neurotransmission is the process by which neurons influence the activity of their target cells via the use of specific signaling molecules known as neurotransmitters. This process involves several steps: first, neurotransmitters are synthesized in presynaptic terminals; then, they are transported into secretory vesicles, released via exocytosis upon neural activity at synaptic and non-synaptic sites, and bound to specific receptors or channels in the target cells such as postsynaptic neurons or myocytes (*1, 2*). Finally, transmission is terminated by degrading enzymes or the transmitter is recycled via reuptake transporter proteins (*3*). Small-molecule neurotransmitters comprise two large classes, 1) the amino acids, such as glutamate, GABA, and glycine (*4*), and 2) the biogenic amines, comprising the monoamines (the decarboxylation and/or hydroxylation products of aromatic amino acids) and acetylcholine. Strictly, acetylcholine is a choline ester bearing a quaternary ammonium group rather than an amine and is often treated as a transmitter class of its own; we here use the term biogenic amines in the inclusive sense of the NLM Medical Subject Headings, which classify acetylcholine together with the monoamines (MeSH D001679), in line with the GPCR classification, in which the muscarinic acetylcholine receptors belong to the class A aminergic receptor family (*5, 6*). Monoamine neurotransmitters consist of serotonin, histamine, the catecholamines epinephrine, norepinephrine, and dopamine, and trace amines. In particular, the trace amines 2-phenylethylamine, tyramine, octopamine and tryptamine owe their name to their very low concentrations in the vertebrate brain, although some of them are known to constitute important neurotransmitters in invertebrate brains, such as octopamine and tyramine, which are major modulators of locomotion and other core functions in insects and mollusks (*7, 8*). Biogenic amines acquired key roles during the evolution of animals, and regulate a wide range of essential neural and physiological functions, among them behavior, cognition, movement, temperature homeostasis, blood pressure and endocrine secretion. However, their presence seems not to be limited to Metazoa; biogenic amines and some of their synthesizing enzymes also exist in plants, fungi, unicellular eukaryotes and even bacteria (*9–15*), suggesting that their roles predate the nervous system. To date, it is not clear how or when during animal evolution biogenic amines acquired their functions in neuronal signaling, and if they are present in all animals or restricted to certain groups. Over the years, different methodologies, including *in vivo* pharmacology, mass spectrometry, HPLC, and immunohistochemistry have hinted at the presence of biogenic amines in non-bilaterian animal groups, including sponges (*16–18*), ctenophores (*19–22*) and cnidarians (*23–39*). Still, it is debated whether these molecules are indeed present and functional in these organisms or if they are products of symbionts or acquired by diet. Moreover, previous phylogenetic analyses found no synthesizing enzymes, transporters or receptors in any of these phyla, concluding that these molecules are likely bilaterian in origin (*40*). Such analyses are based solely on the presence/absence of the classic bilaterian synthetic pathway components across species, which, given the lack of actual protein functional information across the Tree of Life, may lead to a biased, Bilateria-centric view. Two recent studies have challenged it experimentally: trace amines regulate contractions in the sponge *Spongilla lacustris* (*41*), and in the placozoan *Trichoplax adhaerens* they activate GPCRs related to bilaterian melatonin receptors and alter locomotion and body shape (*42*).

To address these questions, we use a multifaceted approach to analyze diverse aspects of biogenic amine signaling in animals. We provide compelling evidence for the presence of diverse biogenic amines in non-bilaterian metazoa via targeted metabolomics and immunochemistry. Phylogenetic analyses of the enzymes and transporters of biogenic-amine signaling, combined with structural modeling, show that the relevant machinery already existed in non-bilaterian animals in a broad-specificity form. Analysis of single-cell expression covariation then places this machinery in specific cell types, both neuronal and non-neuronal, supporting an origin of amine signaling that predates and extends beyond dedicated neural circuits. More specific pathways then emerged via gene duplications in Bilateria. In particular, acetylcholine first emerged as a by-product from carnitine-acetyl shuttling in metabolically active cells, together with muscarinic receptors and nicotinic channels that were expressed in neurons and muscle cells. This suggests that cholinergic interneuron and neuromuscular signaling was already present in the cnidarian–bilaterian ancestor. These results reframe biogenic-amine signaling as a pre-bilaterian property of animal cells, built from promiscuous metabolic enzymes and broad-specificity transporters already present in the last common ancestor of Metazoa, and progressively specialized in bilaterians through repeated duplication and substrate refinement.

## Results

### A broad array of monoamines and acetylcholine are present in sponges, cnidarians, and annelid worms

We tested directly for the presence of biogenic amines in representatives of major animal phyla, and detected a broad array of monoamines including dopamine (DA), histamine (H), adrenaline (epinephrine, Epi), tyramine (TA), 2-phenylethylamine (PEA) and tryptamine (TPA) in extracts of the demosponge *Spongilla lacustris* (Porifera), the freshwater polyp *Hydra vulgaris* and the sea anemone *Nematostella vectensis* (medusozoan and anthozoan Cnidaria respectively), and the marine annelid *Platynereis dumerilii* (Bilateria) (Fig. 1 and Table S1). Both cnidarians were starved before sample preparation (see Methods), as their main food source in the lab (the brine shrimp *Artemia*) would otherwise influence the analysis. Octopamine (OCT) was detected in the annelid and the anemone, and 5-HT in the annelid and the polyp only. Dopamine in the sponge was above the limit of quantification in only one of two replicates (Table S1). L-DOPA and noradrenaline (norepinephrine, NE) were below the limits of quantification for all species (Table S1). Given the occurrence of their immediate downstream metabolites, DA and Epi (Fig. 2A), their absence could be suggestive of their immediate processing, or an alternative biosynthetic pathway (discussed below). Acetylcholine (ACh) was detected in all four species, although in higher concentration in the annelid, whereas acetylcarnitine (ALCAR) was found in all four samples in relatively high concentrations. Note that ACh is expected to naturally co-occur with ALCAR given that carnitine acetyltransferase (CrAT), although it uses L-carnitine as its primary substrate for the synthesis of ALCAR (*43*), can also transfer, to a lesser extent, an acetyl group to choline for ACh synthesis (*44–47*). In contrast, choline acetyltransferase (ChAT) transfers acetyl groups to choline in a highly specific manner to produce ACh (*48, 49*). The ACh:ALCAR ratio was higher in the annelid (≈3:1) than in the sponge (≈1:4), polyp (≈1:60) and anemone (≈1:1,300) (Table S1).

**Figure 1.**
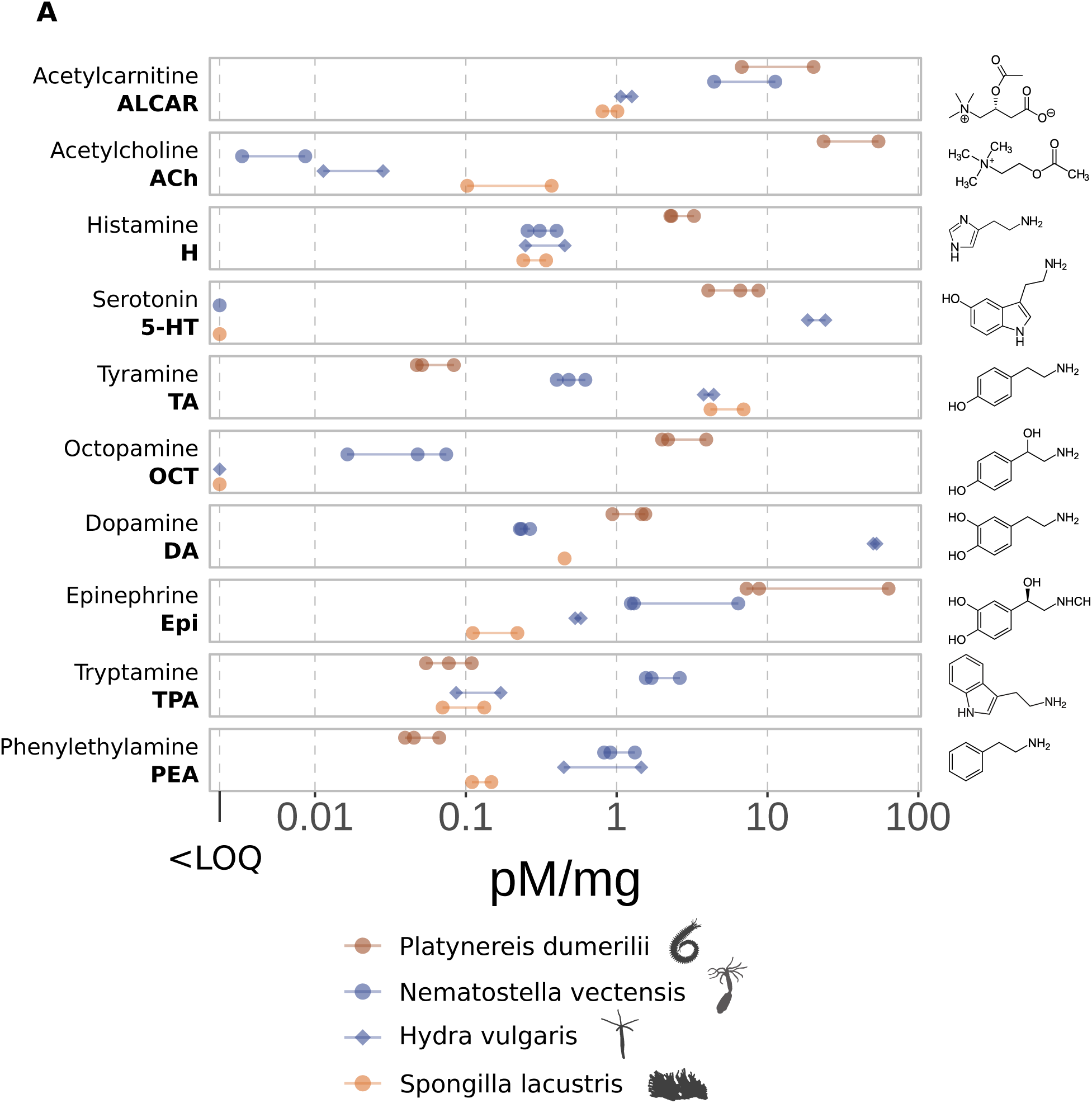
Targeted metabolomics detects biogenic amines in a sponge, two cnidarians, and an annelid. The levels of different biogenic amines (in pM adjusted to mg of sample input) measured in each of the four animal species screened in the metabolomic analysis.

**Figure 2.**
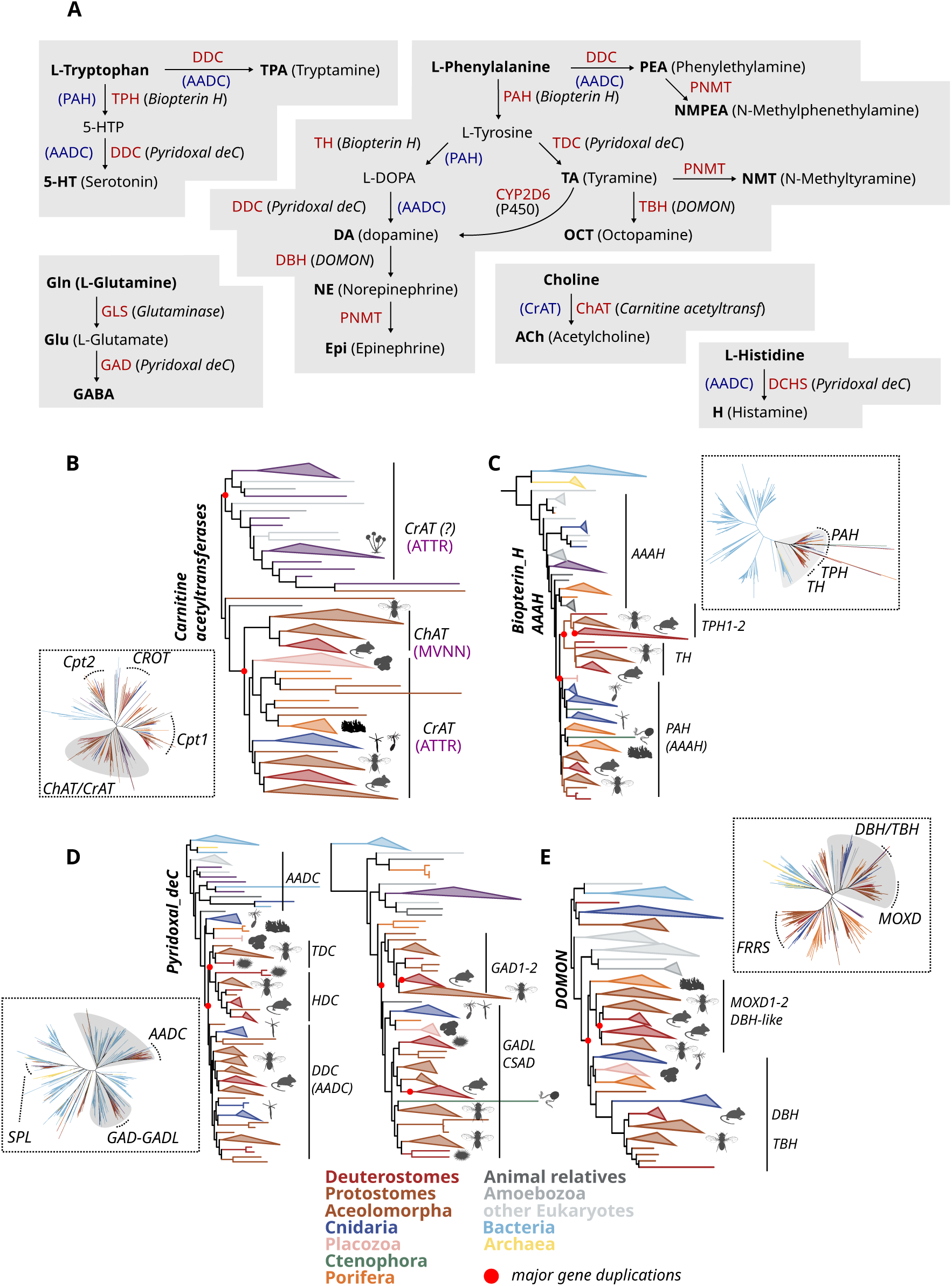
Main biosynthetic pathways of biogenic amines and their evolution. (**A**) Key steps of the main pathways for the production of biogenic amines. The main enzymes that catalyze each reaction are shown in red, alternative enzymes that may promiscuously catalyze the same reaction are shown in blue. The protein family for each enzyme is shown in brackets. (B–E) ML phylogenetic trees for the protein families involved, (**B**) Carnitine acetyltransferases, (**C**) Biopterin­dependent aromatic amino acid hydroxylases, (**D**) Pyridoxal­5’­phosphate­dependent amino acid decarboxylases, and (**E**) DOMON domain­containing β­monooxygenases. An unrooted tree annotated with the main subfamilies for each of the families (those selected for focused analysis in bold) is shown within each panel. The clade which was extracted and re­computed is shown in a rectangular rooted form on the right. The colors representing sequences of different taxonomic origin are shown in the legend below. The red dots on the trees represent the main gene duplication events in each family. In (**B**), the conserved “MVNN” (ChAT­like activity) and “ATTR” (CrAT­like activity) motifs are shown in purple for each main clade.

### Acetylcholine as a by-product of acetylcarnitine production

We set out to explain the aforementioned observations and understand the evolution of neurotransmitter biosynthesis in animals through Maximum Likelihood (ML) molecular phylogenies for each protein family involved in biogenic amine biosynthesis (Fig. 2A). We based our analysis on a broad collection of genomic and transcriptomic data of 44 animals of all five major animal groups, complemented by 24 representative non-animal species (Table S2). ChAT and CrAT both belong to the same family of carnitine transferases (Pfam: PF00755). The phylogenetic tree revealed that ChAT and CrAT derive within the carnitine/choline acetyltransferase clade (Fig. 2B). Bona fide CrATs are found in all bilaterian, cnidarian, and sponge species examined, whereas the ChAT enzymes are only found in Bilateria. More basal sequences co-orthologous to both ChAT and CrAT subclades were detected in many non-animal species, including unicellular animal relatives (*Capsaspora owczarzaki*), Holomycota (fungi and *Fonticula alba*) and other microbial eukaryotic species (see also Fig. S2).

Specific amino acid (aa) residues within the substrate-binding site that differ between ChAT and CrAT have been previously shown to be crucial for carnitine and choline ligand specificity (*50–54*). Directed mutagenesis of a motif of four such residues is sufficient for reversing the catalytic efficiency toward L-carnitine or choline (*51, 55*). We inspected these residues in the multiple sequence alignments (MSAs) across phylogeny (Fig. 2B and Fig. S2) and confirmed that the “MVNN” motif is highly conserved and specific for bilaterian ChAT orthologs, whereas the “ATTR” motif characterizes bilaterian CrATs but is also prevalent in CrAT orthologs in non-bilaterian animals, and in most of the ChAT/CrAT co-orthologs (upper clade in the tree in Fig. 2B). This, together with the fact that all other transferase families have carnitine as their main substrate, strongly suggests that the choline-to-acetylcholine specific biosynthetic activity only evolved (through gene duplication) – and is conserved – in Bilateria.

In an effort to better elucidate the structural and mechanistic aspects of the functional divergence between ChAT and CrAT proteins, we used the Sequence Harmony (SH) method (*56*) to infer key residues related to changes in substrate specificity (see Methods). Interestingly, the most discriminatory residues between the bilaterian ChAT sequences and the other acetyltransferases were found near the enzymatic active center. We mapped them on crystal structures of the human ChAT (PDB: 2FY3), and the murine CrAT (PDB: 2H3P), as well as ColabFold predictions (*57, 58*) of the sponge, anemone and polyp CrAT orthologs. Our analysis suggested that CrATs allow promiscuous reactions through a wide substrate channel (T105, A106; mouse CrAT numbering, positions 201 and 202 of human ChAT in Fig. S3) and large but positively charged active site (R518; position 622 in Fig. S3) (Fig. S3A, B), preferentially allowing the binding of negatively charged substrates such as carnitine. The transition to a choline-specialized enzyme was possible by introducing residues that excluded larger and negatively charged substrates, either directly at the channel entrance (T105D, A106M) or in the active site (R518N) while at the same time favoring the binding of positively charged choline (Fig. S3). Virtually all residues important for CrAT promiscuity are present in the sponge and polyp CrATs, pointing to their potential to use both carnitine and choline as a substrate. The sea anemone orthologs show a diverse combination of residues at these positions, however all favoring CrAT-like activity (Fig. S3C). In extreme cases such as v1g203043, the substrate channel and active site are positively charged, most likely excluding positively charged substrates such as choline (Fig. S3D).

ChAT is therefore a bilaterian specialization, not a bilaterian invention: the catalytic potential for acetylcholine production was already present in CrAT-like enzymes before animals, and the residues necessary for that activity are conserved in sponge and cnidarian orthologs.

### Promiscuous enzymes drive monoamine synthesis in non-bilaterians

Monoamine biosynthesis is dominated by two families, the Biopterin-dependent Hydroxylases (Fig. 2C, Pfam: PF00351), which add hydroxyl groups to their substrates, and the Pyridoxal Decarboxylase family (Fig. 2D, Pfam: PF00282), which decarboxylate aromatic amino acids. Combinations of members of these families constitute the synthesis pathways for most monoamines (5-HT, H, DA, TA, PEA, TPA), complemented by β-hydroxylases of the DOMON family that produce NE and OCT (Fig. 2E, Pfam: PF03351) and methyltransferases of the NNMT/PNMT/TEMT family (Fig. S1A, Pfam: PF01234) that produce Epi and dimethyltryptamine (DMT). The ML trees for all those families revealed the same general evolutionary trend: widespread taxonomic distribution of broad substrate specificity, limited distribution of narrow specificity; the more promiscuous family members are conserved across large evolutionary distances, whereas enzymes with single or fewer substrates are found in more restricted clades only. The latter holds true for example for tyrosine hydroxylase (TH, EC 1.14.16.2) for the production of L-DOPA and tryptophan hydroxylase (TPH, EC 1.14.16.4) for 5-HTP (Fig. 2C); Tyramine-β-hydroxylase and Dopamine-β-hydroxylase (TβH and DβH, EC 1.14.17.1) for OCT and NE (Fig. 2E), and tyrosine and histamine decarboxylases (TDC, EC 4.1.1.25, and HDC, EC 4.1.1.22) for the production of TA and H (Fig. 2D, left), and even glutamate decarboxylases 1 and 2 (GAD1/2, EC 4.1.1.15), part of the Pyridoxal Decarboxylase family, which catalyze the synthesis of GABA (Fig. 2D, right). These enzymes represent specialized Bilateria-specific duplicates. In contrast, homologous enzymes that we found conserved across non-bilaterian metazoans represented enzyme family members reported to be promiscuous, i.e. driving the production of several monoamines (Fig. 2C–E). Cnidarians, *Trichoplax*, sponges, and ctenophores (long branch nested within sponge sequences in Fig. 2C) encode a protein referred to as phenylalanine hydroxylase (PAH, EC 1.14.16.1) due to the production of L-tyrosine from L-phenylalanine, but it is also known to catalyze the synthesis of 5-HTP from tryptophan, substituting the function of TPH for serotonin synthesis (*14, 59, 60*). Similarly, non-bilaterian metazoans encode proteins related to DOPA decarboxylase (DDC, EC 4.1.1.28), often referred to more generally as aromatic amino acid decarboxylase (AADC) (Fig. 2D, left). This enzyme has the striking ability to decarboxylate L-DOPA, 5-HTP, L-histidine, L-phenylalanine, L-tryptophan and L-tyrosine to produce DA, 5-HT, H, PEA, TPA and TA respectively (*61–65*). Notably, in sponges monoamine synthesis may not happen through this ortholog at all. In *Spongilla lacustris*, tyramine, 2-phenylethylamine and tryptamine production has recently been attributed to PLP-dependent decarboxylases outside the canonical AADC clade (*41*), broadening the set of enzymes able to supply trace amines in non-bilaterians.

The analysis and interpretation of the evolution of the biosynthetic enzymes of biogenic amines suggest that the first animals were already capable of generating a vast array of monoamines via rather unspecific members of the Biopterin Hydroxylase, Pyridoxal Decarboxylase, DOMON β-Hydroxylases and Methyltransferase families. Gene duplications then led to the emergence of more specific enzymes in the bilaterian lineage. The cnidarian-specific expansions of the same families suggest that comparable specialization may have occurred independently in this lineage; the substrate preferences of these paralogs, however, remain elusive and await direct biochemical testing.

### Promiscuous transporters for acetylcholine and monoamines in non-bilaterian metazoans

The Major Facilitator Superfamily (MFS, Pfam: PF07690) is the largest secondary active transporter superfamily, and includes at least 16 solute carrier (SLC) families (*66*). Characterized members transport a remarkably broad range of substrates, including diverse neurotransmitters. The Vesicular Acetylcholine Transporter (VAChT, *Slc18a3*), the vesicular monoamine (VMAT/*Slc18a1-2*) and the vesicular polyamine (VPAT/*Slc18b1*) transporters are all members of the SLC18 family of MFS transporters (Fig. 3A). A different MFS family, the SLC22, includes many broad-specificity transporters (at least 23 genes in human), with substrates ranging from organic anions and cations, carnitine, choline and creatinine, but also ACh, DA, NE, Epi among other molecules (*67–69*). Some SLC22 members have been shown to be enriched in cholinergic neurons, localized in the synaptic vesicles in the presynaptic terminals of motor neurons and the NMJ (*70*). To understand the evolution of neurotransmission-related vesicular transport, we detected all MFS proteins in our species, split them into families using a similarity-based clustering approach (see Methods), and analyzed phylogenetically the SLC18 and SLC22 transporter families (Fig. 3A).

**Figure 3.**
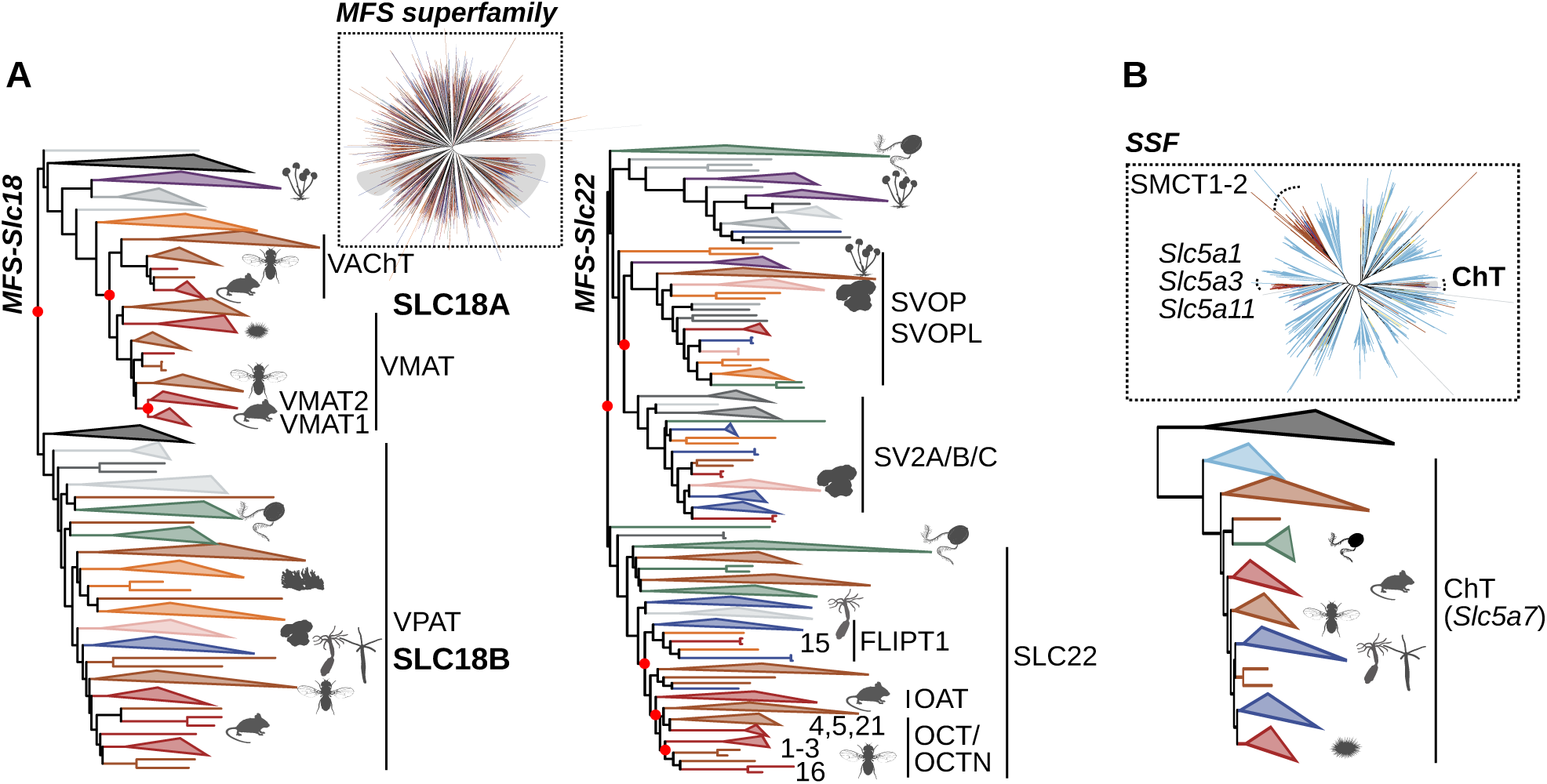
The evolution of biogenic amine vesicular and reuptake transporters. (**A**) Phylogenies of the Major Facilitator Superfamily and the SLC18 (left) and SLC22 (right), and (**B**) the SLC5 transporter family. (**A**) The VAChT and VMAT clades (left) are paralogous “sister clades”, deriving from a gene duplication that happened in the bilaterian stem lineage, as inferred by the presence of deuterostome (dark red) and protostome (brown) sequences in both clades. Non­ bilaterian co­orthologs to both VAChT and VMAT exist in sponges (orange), fungi (purple), amoebozoa (dark gray) and other eukaryotes (light gray), with no functionally characterized member. The VPAT clade is widely present in all metazoa and non­metazoan eukaryotes. The SLC22 family phylogeny (right) is reconstructed with a reduced representative taxon sampling (see Methods) due to its large size. It is also widespread across animals and paralogous to both synaptic vesicular (SV2) and synaptic vesicle 2­related protein (SVOP) clades. (**B**) The high affinity choline transporter (ChT) is part of the SSF family (SLC5) of transporters, which includes the sodium­coupled monocarboxylate transporters (SMCT­1/2), the sodium/myo­inositol (Slc5a3/11) transporters, and sodium/glucose co­transporters. All colors are as in Fig. 2.

In line with previous observations, the SLC18 tree showed that SLC18A (VAChT and VMATs) and SLC18B (VPAT) form two paralogous eukaryotic clades (Fig. 3A, left), implying that early eukaryotes had one ancestral SLC18 vesicular transporter of possibly broad specificity which duplicated and diverged into A and B subfamilies. A subsequent duplication of the SLC18A gene in the bilaterian stem separated the VAChT and VMAT subclades. This suggests that the differential and more specific transport of ACh and monoamines into (synaptic) vesicles became possible only in Bilateria. Notably, VAChT and VMAT co-orthologs (i.e., basal SLC18A members) are found in some sponges (*Sycon ciliatum* and *Oscarella lobularis*), the cnidarian *Cladocora caespitosa*, animal relatives (*Capsaspora owczarzaki and Sphaeroforma arctica)*, all seven fungal species in our dataset, *Acanthamoeba castellanii* (Amoebozoa), but also some algae across our taxon sampling (*Emiliania huxleyi, Thalassiosira pseudonana, and Guillardia theta*), suggesting that the A clade was lost in many animal and non-animal lineages, including the model cnidarians and most of the sponges investigated here. Within the SLC18B clade, the VPAT/*Slc18b1* orthologs are more widespread, found in all animal groups and other eukaryotes, except for *A. castellanii* and fungi. VPAT is able to load a range of amine substrates into vesicles, including acetylcholine, monoamines such as histamine and serotonin, and polyamines such as spermidine and spermine (*71, 72*).

The SLC22 family tree (Fig. 3A, right; single-cell expression in Figs. S9, S10) shows that the diversification of the different SLC22 subfamilies of transporters (the organic anion transporters, OATs; the organic cation transporters, OCTs; and the carnitine/organic cation transporters, OCTNs) occurred within Bilateria. Co-orthologs are, however, detected in cnidarians, ctenophores and sponges, but also plants (called ‘organic cation transporters’ in *A. thaliana*). Most representatives of this family are known to be promiscuous transporters with multiple structurally different substrates, ranging from polyamines, monoamines, acetylcholine, carnitine, choline, nucleotides, urates, and different hormones (*73–80*). The phylogeny also illustrates the close relatedness of the SLC22 family with the synaptic vesicle glycoprotein 2 (SV2) and SV2-related protein (SVOP), both widespread in animals but, interestingly, also in their unicellular relatives and fungi (dark gray and purple). While the functions of SV2 and SVOP proteins have not been completely elucidated, it is known that these proteins are located in synaptic neurotransmitter-containing vesicles, and both are involved in the regulated secretion of neurotransmitters from presynaptic neurons (*81*). In humans, the SV2A protein is ubiquitously expressed in the brain, neuroendocrine cells and at the NMJ (*82–84*). In *Spongilla*, SV2/SVOP-related MFS transporters are co-expressed with the biosynthetic enzymes in secretory cells and support monoamine-dependent contractility (*41*), so promiscuous vesicular loading in non-bilaterians seems to extend beyond the SLC18B/VPAT clade we emphasize here. Reuptake follows the same logic: the high-affinity choline transporter (ChT), which recycles choline at bilaterian cholinergic terminals, is a bilaterian-specific member of the broadly conserved SLC5 transporter family (Fig. 3B).

Together, these data suggest that unspecific vesicular loading of monoamines and acetylcholine was possible long before the emergence of Bilateria, and, at least in Metazoa, mostly carried out by broad-specificity transporters, including VPAT clade members and SV2/SVOP-related transporters. As shown for the synthesizing enzymes, more promiscuous transporters evolved first, followed by more substrate-specific transporters such as VAChT and VMAT in Bilateria (and possibly also in cnidarians and other groups, as suggested by the high number of species-specific duplicates of unknown substrate specificity). Across all three layers of the cholinergic and monoaminergic machinery we have analyzed so far (biosynthesis, vesicular loading and reuptake), the same pattern emerges: broad-specificity ancestors retained across animals, narrow-specificity bilaterian gene expansions.

### Bona fide mAChR and nAChR outside bilaterians

For signals to be relevant, they need to be specifically received. We explored whether receptors for biogenic amines exist in non-bilaterian metazoa and beyond, by reconstructing and analyzing phylogenies of the protein superfamilies of the two major types of neurotransmitter receptors, the metabotropic G protein-coupled receptors (Rhodopsin family of GPCRs, Pfam: PF00001) and the ionotropic Cys-loop ligand-gated ion channels (Pfam: PF02931 and PF02932).

#### Medusozoa encode mAChRs

Different GPCR analyses using alternative phylogenetic algorithms and taxon samplings suggested that the classic metabotropic monoamine, acetylcholine, and histamine receptors are more closely related (Fig. 4A), to the exclusion of adenosine, cannabinoid, melatonin receptors, and peptidergic GPCRs. Our tree analyses support the previous observation that the characterized monoamine receptors (apart from histamine) are found only in Bilateria, with no orthologous genes in non-bilaterian species. This does not preclude lineage-specific monoamine receptors in non-bilaterians, as ligand specificity can shift with a few substitutions in the binding pocket; indeed, *Trichoplax adhaerens* orthologs of bilaterian melatonin receptors respond to non-acetylated monoamines rather than N-acetylated ligands (*42*). In contrast to the dedicated monoamine receptors, we detected cnidarian sequences directly orthologous to the bilaterian muscarinic acetylcholine receptors (mAChRs), as well as cnidarian, placozoan, and ctenophore sequences related to the bilaterian H2 receptors (Fig. 4A). The bona fide mAChR orthologs were identified in medusozoan cnidarians, and have been lost in anthozoans. The residues that best discriminate the cnidarian–bilaterian mAChR clade from the other GPCRs lie in the ligand-binding pocket (Fig. 4C). Focusing on the *H. vulgaris* mAChR orthologs, repeated gene duplications resulted in various copies within two main subclades (A and B, Fig. 4B). A sequence comparison of the orthosteric ACh binding site based on the human M5 mAChR (*85*) revealed 1) complete conservation of all residues in this site within the bilaterian mAChRs, and 2) complete conservation in two *H. vulgaris* receptors within the A subclade (Fig. 4B, XP_012561685.1 and XP_012561731.1, and the matching transcriptome sequence t21045aep). All *H. vulgaris* receptors of the B subclade had three or more amino acid changes in the orthosteric site (Fig. 4B).

**Figure 4.**
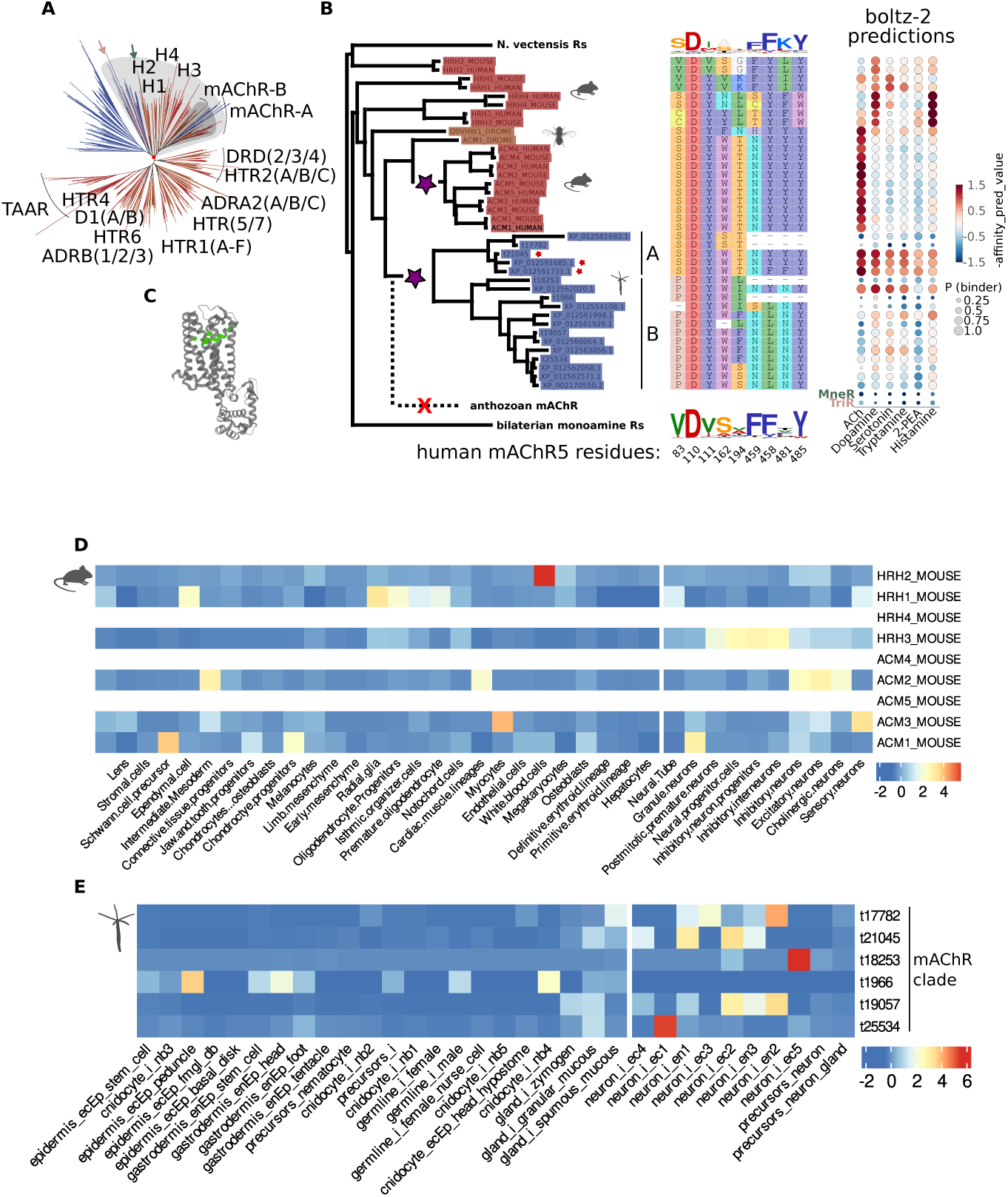
Metabotropic ACh­sensitive receptors in *Hydra vulgaris*. (**A**) GPCR phylogeny of the H­ACh­monoamine clade. The parts of the tree have been annotated based on the characterized bilaterian receptors. Whereas the monoamine receptors monophyletic clade is Bilateria­specific, medusozoan sequences are found to be nested within the AChRs and orthologous to mAChRs (in blue). Other cnidarian, ctenophore (MneR, green arrow) and placozoan (TriR, pink arrow) sequences are found closer to the H2 bilaterian receptors or more distantly. The light gray­shaded area represents the clade that was extracted and re­computed, and the darker gray the part that is shown in (**B**). (**B**) Phylogenetic relationships of the human, mouse, fly and *H. vulgaris* members of the H­ACh GPCR clade, and the amino acid residues in the ligand orthosteric binding site, according to the human M5 mAChR (*85*). The sequence conservation logos of the related *N. vectensis*, and the monoamine receptor sequences were created using the WebLogo server (*134*). The *H. vulgaris* sequences with an orthosteric site identical to that of the bilaterian mAChRs (ACh­rec1, ACh­rec2 and the matching transcriptome sequence t21045aep) are indicated with a red star. Purple stars indicate the single ancestral mAChR of deuterostomes and of medusozoans, and the red cross its loss in anthozoans. (**C**) The top ranking (according to Sequence Harmony (SH) method (*56*)) discriminatory amino acids between the cnidarian–bilaterian mAChR clade (gray shaded) and all the other GPCRs in (**A**) are in the ligand binding pocket (indicated in green on the M5 mAChR structure). (B, right) Boltz­2 structural binding­affinity predictions for each receptor against six candidate ligands (acetylcholine, dopamine, serotonin, tryptamine, 2­phenylethylamine and histamine); dot size indicates the predicted binding probability and color the predicted affinity (−affinity_pred_value; higher values indicate stronger binding). Receptors with experimentally validated specificities (bilaterian mAChRs and HRH2–4) recover their known ligands; the *Hydra* receptors with a conserved orthosteric site (red stars; ACh­rec1, ACh­rec2 and the matching transcriptome sequence t21045aep) show the strongest acetylcholine predictions in the *Hydra* set; the other *Hydra* receptors show weaker predictions, with the exception of XP_012562020.1; the ctenophore (MneR) and placozoan (TriR) receptors show low predictions for all ligands. (D,E) Normalized (z­score) single­cell expression of (D) mouse H­ ACh­clade receptors and (E) *H. vulgaris* mAChR­clade receptors across cell types. Mouse H3 and mAChRs and the *H. vulgaris* receptors are predominantly expressed in neuronal cell types, mouse H1 and H2 are not; the broad mAChR expression reflects the wide effects of cholinergic signaling. The transcriptome­derived *H. vulgaris* receptor t21045aep, which has a totally conserved ligand­binding site, is most highly expressed in ectodermal and endodermal neuron types (ec2, ec4, en1, en3).

To assess the predicted ligand specificity of the *H. vulgaris* mAChR orthologs we used the structural binding-affinity prediction tool Boltz-2 (*86*) to score receptors against six candidate ligands: acetylcholine, histamine, dopamine, serotonin, tryptamine and 2-phenylethylamine (Fig. 4B). The method correctly recovered the known specificities of the bilaterian positive controls, namely acetylcholine for the human, mouse and *Drosophila* mAChRs, histamine for HRH2–4 (though not for HRH1), serotonin for 5-HT2A–C and dopamine for DRD2–4 (Fig. S15; Table S4), and gave uniformly low predictions for the ctenophore and placozoan receptors MneR and TriR (arrows in Fig. 4A), supporting the reliability of the predictions before we examined the *Hydra* receptors. Among the *H. vulgaris* mAChR orthologs, the two with a fully conserved orthosteric site, ACh-rec1 (XP_012561685.1) and ACh-rec2 (XP_012561731.1), gave the strongest acetylcholine predictions of any *Hydra* receptor (probability ∼0.997, comparable to the bilaterian mAChRs). The other *H. vulgaris* receptors, with substitutions in the orthosteric site or incomplete sequences, gave weaker predictions for acetylcholine (probability 0.06–0.74, mostly with positive affinity values, i.e. predicted IC50 above 1 µM), with the single exception of XP_012562020.1 (probability 0.975). The transcriptome sequence t21045aep, which corresponds to ACh-rec1/2 and shows predominantly neuronal expression (Fig. 4E), shared the same high acetylcholine binding profile. ACh-rec1 and ACh-rec2 also gave intermediate-to-high binding probabilities for the other ligands (0.83–0.94), with acetylcholine still the most favorable in each case. Because Boltz-2 scores the structural compatibility of the binding pocket rather than receptor activation, these predictions remain to be tested experimentally. Phylogeny, orthosteric-site conservation and structural binding prediction are therefore in agreement: medusozoan cnidarians possess muscarinic acetylcholine receptor orthologs, with ACh-rec1 and ACh-rec2 the most likely acetylcholine-binding receptors in *H. vulgaris*.

We looked at the single-cell transcriptomic profiles of the receptors in whole-body mouse (*87*) and *H. vulgaris* (*88*) data. It is worth noting that the mouse data show that the more basal H2 (related receptors found in Cnidaria, Placozoa, Ctenophora, see also Fig. 4A) and H1 receptors are not expressed in neurons, in contrast to H3 and mAChRs which are more closely related (Fig. 4D). *Hydra* mAChR orthologs are predominantly expressed in neuronal cells, and secondarily in gland cells (Fig. 4E). Notably, transcript t21045aep, which corresponds to ACh-rec1/2 and was the only *Hydra* transcript with the same fully conserved orthosteric site as the bilaterian mAChRs, is most highly expressed in specific ectodermal and endodermal neuronal cell types and at lower levels in other neuronal and non-neuronal cells (Fig. 4B, E).

#### A nAChR channel in muscle and neurons in the ancestor of Cnidaria and Bilateria

The nicotinic acetylcholine receptors (nAChRs) belong to the Cys-loop receptors, and convey fast neurotransmission in bilaterians. The functional channels are homo- or hetero-pentameric, composed of subunits arranged around a central transmembrane pore. Our phylogenetic analysis of the superfamily suggests that an ancient duplication separated the subunits of anion-selective (predominantly GABA and glycine, but also glutamate and monoamines) and cation-selective (ACh, 5-HT) channels (Fig. 5A). In agreement with previous observations (*89*), we detected orthologs of nAChRs in both anthozoan and medusozoan cnidarians (Fig. 5A and B), indicating that the emergence of the ionotropic nicotinic AChRs occurred in the cnidarian–bilaterian common ancestor (CBA). The ancestral CBA nAChR was homo-pentameric, assembled from five copies of one subunit from which all cnidarian and bilaterian nAChR subunits originate (arrow in Fig. 5B). The cnidarian nAChR subunits form a monophyletic clade, sister to the bilaterian nAChR clade, indicating that the bilaterian and cnidarian subunits expanded and diversified independently. The whole nAChR subunit clade is sister to the 5-HT3Rs which were subsequently lost in cnidarians but retained in Bilateria. The subunit composition of the assembled pentameric channels differs across cells and tissues, which defines distinct subtypes (e.g. muscle-type, ganglion-type) with distinct functional properties. For instance, in the vertebrate NMJ, which is cholinergic, the postsynaptic muscle cells receive the signal via nAChRs composed of α1, β, γ, δ and ε subunits (marked on tree in Fig. 5B). While both glutamatergic and cholinergic NMJ synapses are known in protostomes (arthropods use Glu, nematodes ACh) (*90*), it is not clear what neurotransmitter non-bilaterians use for neuron–muscle signaling. Given the existence of nAChR orthologs in Cnidaria, we reasoned that if they use ACh for neuromuscular signaling, similarly to vertebrates, they may also have different subunit compositions in muscles and neurons. Initially, we confirmed that in mouse, muscle-type subunits (apart from ε which had no detected expression) are specifically expressed in myocytes, in contrast to other nAChR and 5-HT3R, and other Cys-loop and iGluR subunits which are mostly expressed in neurons (Fig. 5C and Fig. S4). Their expression is strongly correlated with that of Rapsyn, an important scaffolding protein essential for NMJ formation (*91*). In contrast to *D. melanogaster*, which has a glutamatergic NMJ and does not express nAChR subunits in muscles but rather in motor neurons, while iGluRs are expressed in muscle cells instead (Fig. S5), we detected muscle-type subunits both in muscle-associated epidermal cell types in *H. vulgaris* (Fig. 5D and Fig. S6), as well as in the tentacle retractor muscle of *N. vectensis* (Fig. S7), also co-expressed with cnidarian Rapsyn-related genes. Strikingly, we found those muscle-expressed subunits in the two cnidarians to group together in a clade in the tree, suggesting that ACh-sensitive muscle cells were already present in the cnidarian ancestor. These results point to differential cholinergic neuromuscular and inter-neuronal signaling in Cnidaria, with subtypes of distinct subunit composition functioning in the different sites, as in vertebrates. Taking into account the expression profiles in both the bilaterian and cnidarian clades of the tree, we may infer that the CBA had one homo-pentameric channel expressed both in neuronal and muscle cells. Cholinergic neuromuscular signaling is therefore not a bilaterian innovation. Muscle-cell-expressed nicotinic channels co-expressed with Rapsyn were already present in the cnidarian–bilaterian common ancestor; bilaterians inherited this arrangement and specialized the subunits, while arthropods such as *Drosophila* secondarily switched to glutamatergic neuromuscular transmission (Fig. S5). Together with the muscarinic receptor orthologs in cnidarians, these data place the core components of the cholinergic system in the cnidarian– bilaterian ancestor.

**Figure 5.**
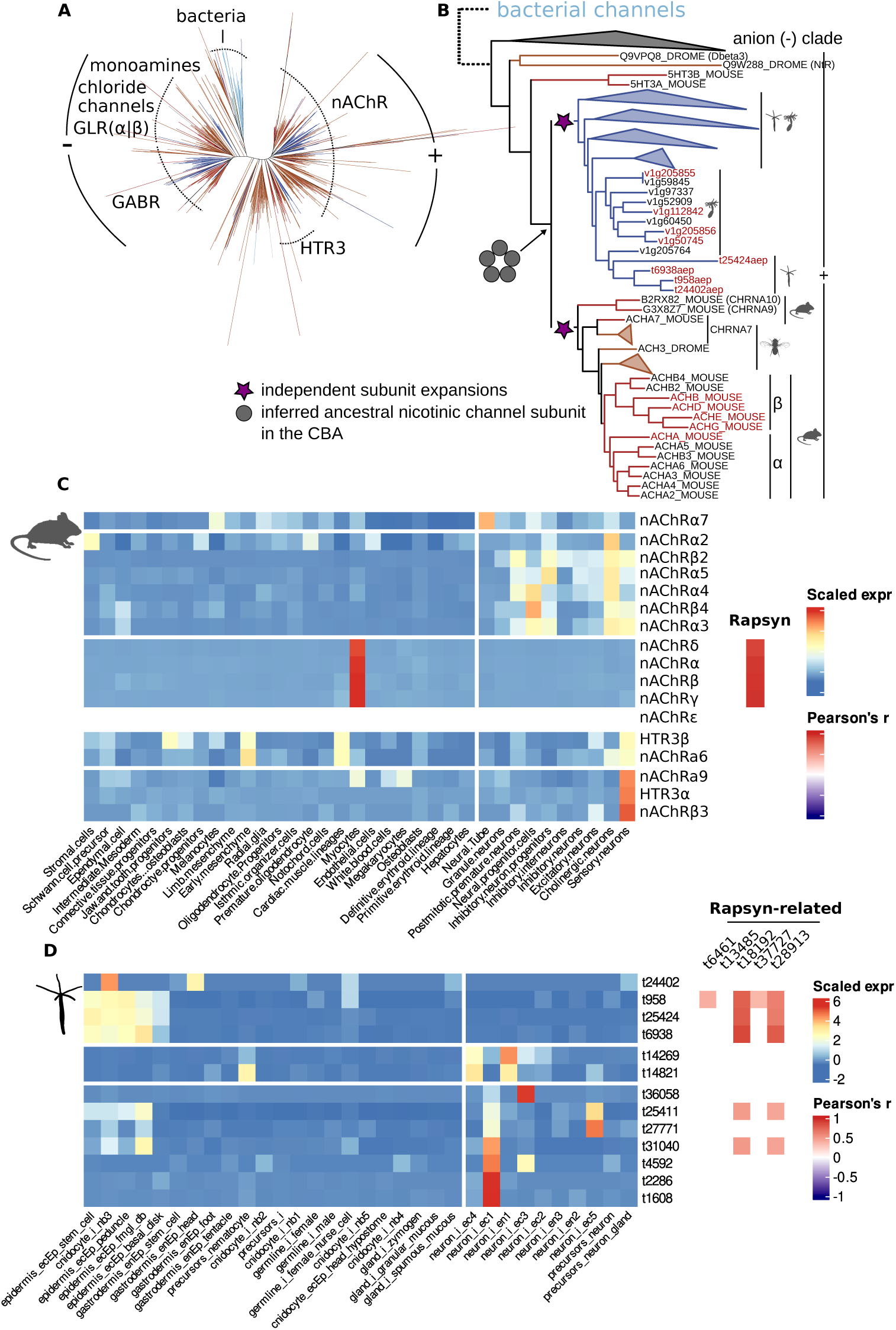
Cnidarian nicotinic ACh receptor channels in neurons and muscles. (**A**) The phylogenetic tree of the Cys­ loop ligand­gated ion channel subunits superfamily shows that anion (GABA­, glycine­, glutamate­, and monoamine­ activated) and cation (5­HT and ACh­activated) channels are composed of subunits that form two phylogenetically distinct clades of potential bacterial origin. (B) The relationships within the cationic clade suggest a common origin of the 5­HT and ACh­sensitive channels from a homo­pentameric channel of unknown sensitivity in the cnidarian–bilaterian ancestor (CBA). Cnidaria probably lost 5­HT channels, but maintained nAChR­related channels, and expanded their subunits in parallel to the bilaterian subunits. (C) In mouse, the nAChR subunit expression is suggestive of the channels’ arrangement as the main mediators of fast excitatory synaptic transmission in ganglionic neurons and the neuromuscular junction (NMJ). Their expression in myocytes is correlated with that of Rapsyn (Pearson’s r on the right). (**D**) Similarly in *H. vulgaris*, distinct subunits are expressed in neuron cell types, and in epidermis (muscle) and cnidocyte cell types, in analogy to the bilaterian members, suggesting a role of nicotinic cholinergic signaling both in cnidarian neurons and muscles, pushing back the origin of the system to the CBA, after which it further specialized and diversified in parallel in the two sister lineages. In (D), the subunits expressed in epidermal (muscle) cell types are correlated with Rapsyn-related genes (Pearson’s r on the right).

### Evidence for neuromuscular cholinergic signaling in *Nematostella vectensis*

The direct detection of ACh in cnidarians, as well as the detection of orthologous muscular nAChR subunits in *H. vulgaris* and *N. vectensis*, prompted us to perform immunochemistry using commercial anti-ACh antibodies, to visualize the cells producing it (Fig. 6). We prepared explants from the body wall of *N. vectensis* and stained them for ACh (see Methods). We found numerous flask-shaped ACh-immunopositive cells, so far uncharacterized, flanking the parietal muscle at the base of each of the eight mesendodermal mesenteries (Fig. 6B,C). We also detected ACh-immunopositive cells with neurite-like bipolar extensions in circumferential direction in the gastrodermal body wall between the mesenteries, perpendicular to the parietal muscle (Fig. 6D,E). These data further support that ACh can be detected in neuronal and non-neuronal cells specifically associated with muscles of the sea anemone, suggestive of a vertebrate-like role of ACh in *N. vectensis* neuromuscular signaling.

**Figure 6.**
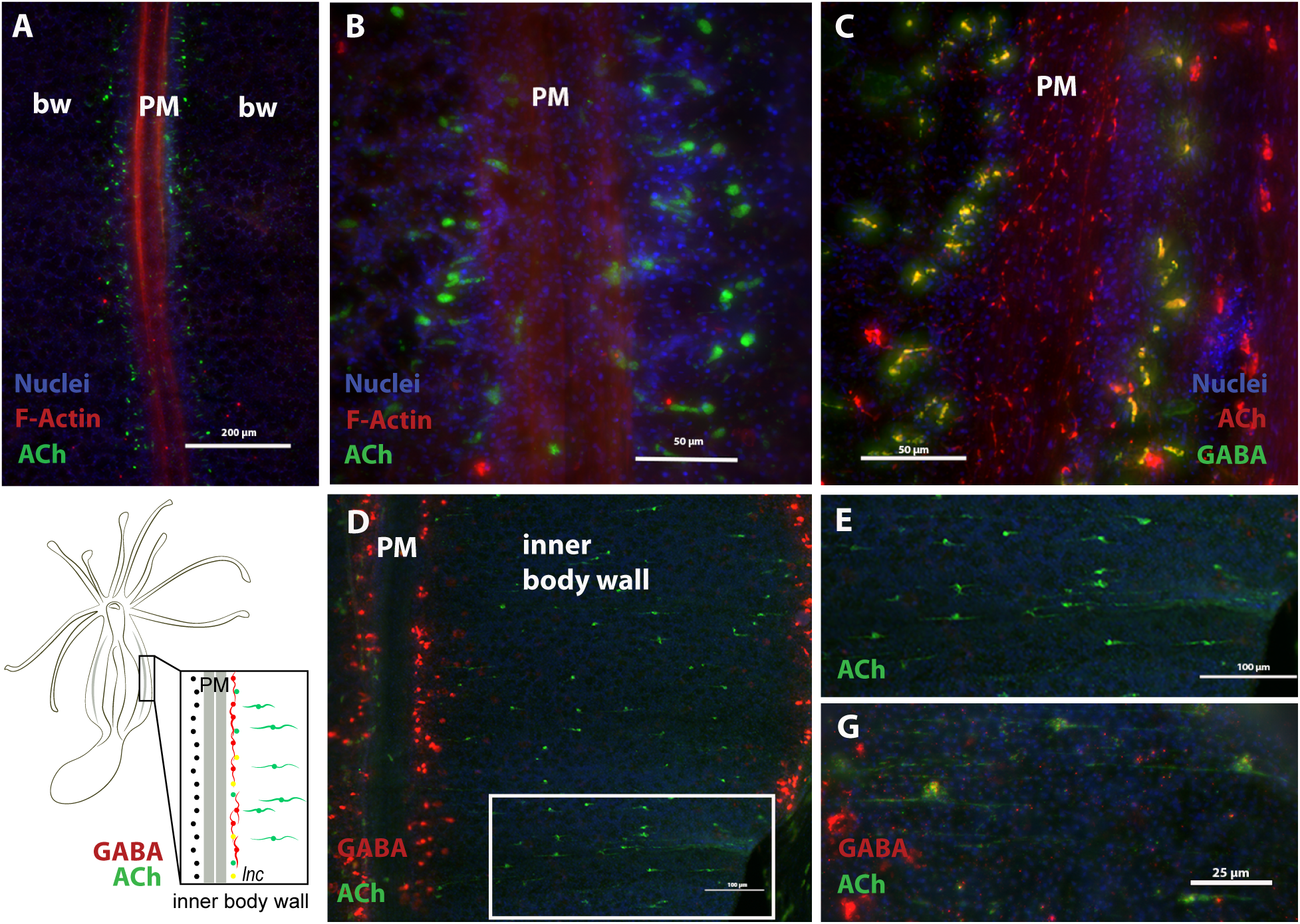
Distribution of acetylcholine immunoreactivity in the body wall of the sea anemone *Nematostella vectensis*. (A) Schematic overview of the body wall sections shown in B–E. **(B,C)** Immunohistochemistry for ACh and phalloidin to highlight the parietal muscle embedded within the body wall. Diverse cell morphologies are found within the area of the longitudinal nerve tracts that run along either side of the parietal muscle. (D,E) ACh immunohistochemistry highlighting cells with a neuron­like morphology running perpendicular to the parietal muscle within the inner body wall epithelia. PM: parietal muscle; lnc: longitudinal nerve cord; bw: body wall. Scale bars: B,D, 200 µm; C,E, 50 µm.

### Candidate cell types releasing acetylcholine or monoamines

The specific distribution and possible coregulation of transcripts in single-cell data is an important source of information to identify candidate cell types producing and releasing acetylcholine or monoamines. To exemplify this approach, we confirmed the specific co-expression of *ChAT, VAChT* and *ChT* genes in cholinergic neurons in mouse (Fig. 7A). ChAT and VAChT expression appears broader in fly cell types (headless dataset), but still correlated and entirely specific for neurons (Fig. S8A). Regarding monoamines, the mouse data revealed exclusively neuronal expression of *TH* for dopamine synthesis. In contrast, two *TPH* paralogs (that specifically synthesize the serotonin precursor 5-HTP) are also expressed in non-neuronal tissue, as is *AADC/DDC,* consistent with broader serotonin production. Interestingly, the strong *AADC/DDC* expression in cardiac muscle correlates with strong co-expression of *PNMT,* demarcating adrenergic muscle lineages (*92*). In *Drosophila* we observe an opposite specificity: strictly neuronal *TPH* expression (in ventral nerve cord and gustatory receptor neurons) versus non-neuronal *TH* expression (e.g., in epithelial cells) correlate with *AADC*/*DDC*, consistent with neuronal serotonin and non-neuronal dopamine production (Fig. S8A).

**Figure 7.**
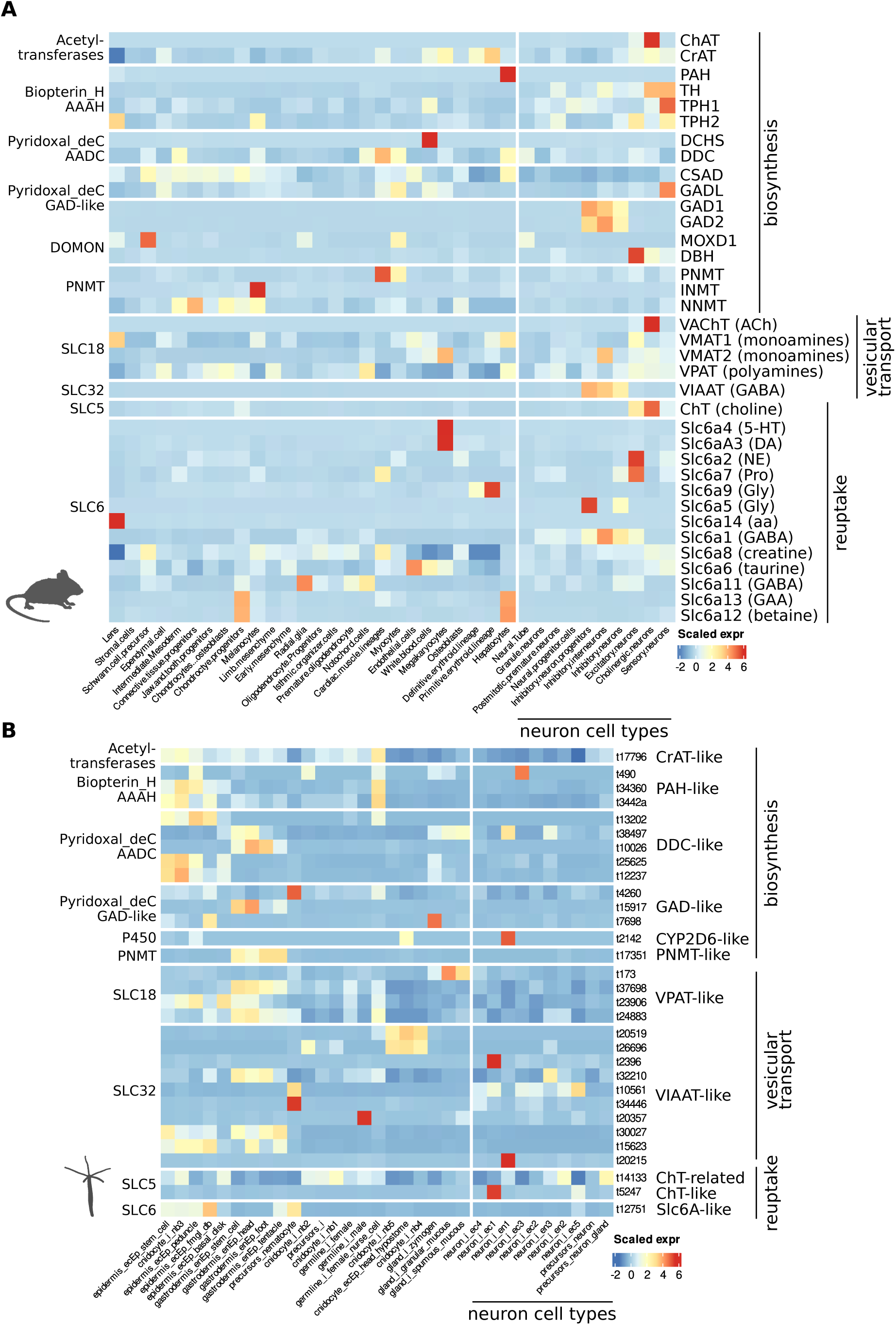
Single­cell expression of biosynthesis enzymes, vesicular, and reuptake transporter genes in mouse and *H.* *vulgaris*. Heatmap of the normalized expression values of the genes within each family for mouse (**A**) and *H. vulgaris* (**B**). *Hydra lacks DOMON­containing proteins* related to the MOXD­DβH clade (see also Fig. 2), and its P450 CYP2D6 related gene is included.

Next, we analyzed single-cell expression data for *Hydra* and *Nematostella* to identify candidate cell types synthesizing ACh or monoamines. In *Hydra*, we observed broad upregulation of *AAAH* and *AADC* paralogs in gastrodermis and epidermis (Fig. 7B) and single neuron types showing strong and specific upregulation of single paralogs of either *AAAH* or *AADC* (not both). In *Nematostella*, one paralog each of AAAH and of AADC is broadly expressed, whereas a second paralog of *AADC* is exclusively expressed in one neuron type (Fig. S8B). This would allow for the production of 5-HT and DA (requiring both enzymes) and TA, TPA, H and PEA (requiring *AADC* only) from L-tryptophan, L-phenylalanine, L-tyrosine or L-histidine (Fig. 2A), which are indeed detected in both species (with the exception of 5-HT in *Nematostella*). Notably, DβH-like monooxygenases were only found expressed in *Nematostella*, in line with the detected presence of OCT in *Nematostella* only (and higher Epi content in *Nematostella*). We also noted that one of the *Hydra AADC*+ neurons (neuron_i_en1) specifically co-expresses an ortholog of *CYP2D*, which provides an alternative route for DA synthesis from TA (*93, 94*), which makes it a candidate DA producing cell type.

### Single-cell co-expression of amine-metabolism genes in *Hydra, Nematostella* and *Mnemiopsis*

To explore choline and monoamine metabolism more broadly in non-bilaterian metazoans, we identified, for each species, the set of genes co-expressed with the seed genes of eight biosynthesis and transport families, including AAAH and AADC (see Methods), across cell types, and intersected this set with the GO:0009308 annotation ("amine metabolic process"). For *Hydra, Nematostella* and *Mnemiopsis* we obtained species-specific co-expressed gene sets, which we call coregulons (Fig. 8; Fig. S11). In *Hydra*, the combined coregulon was distributed across gastrodermal, epidermal, germline and cnidocyte cell types, with a few members (PAH, AMD1, AFMID, a NAPE-PLD paralog and an SLC5 transporter) peaking in neuron types (Fig. 8); in *Nematostella*, the combined coregulon spanned gland, muscle and epidermal cell types, and about a third of its members, including CRAT, two SLC18B1 paralogs and five SNF transporters, peaked in neuron types (Fig. S11A); in *Mnemiopsis*, the coregulon was distributed across non-neuronal cell types (Fig. S11B).

**Figure 8.**
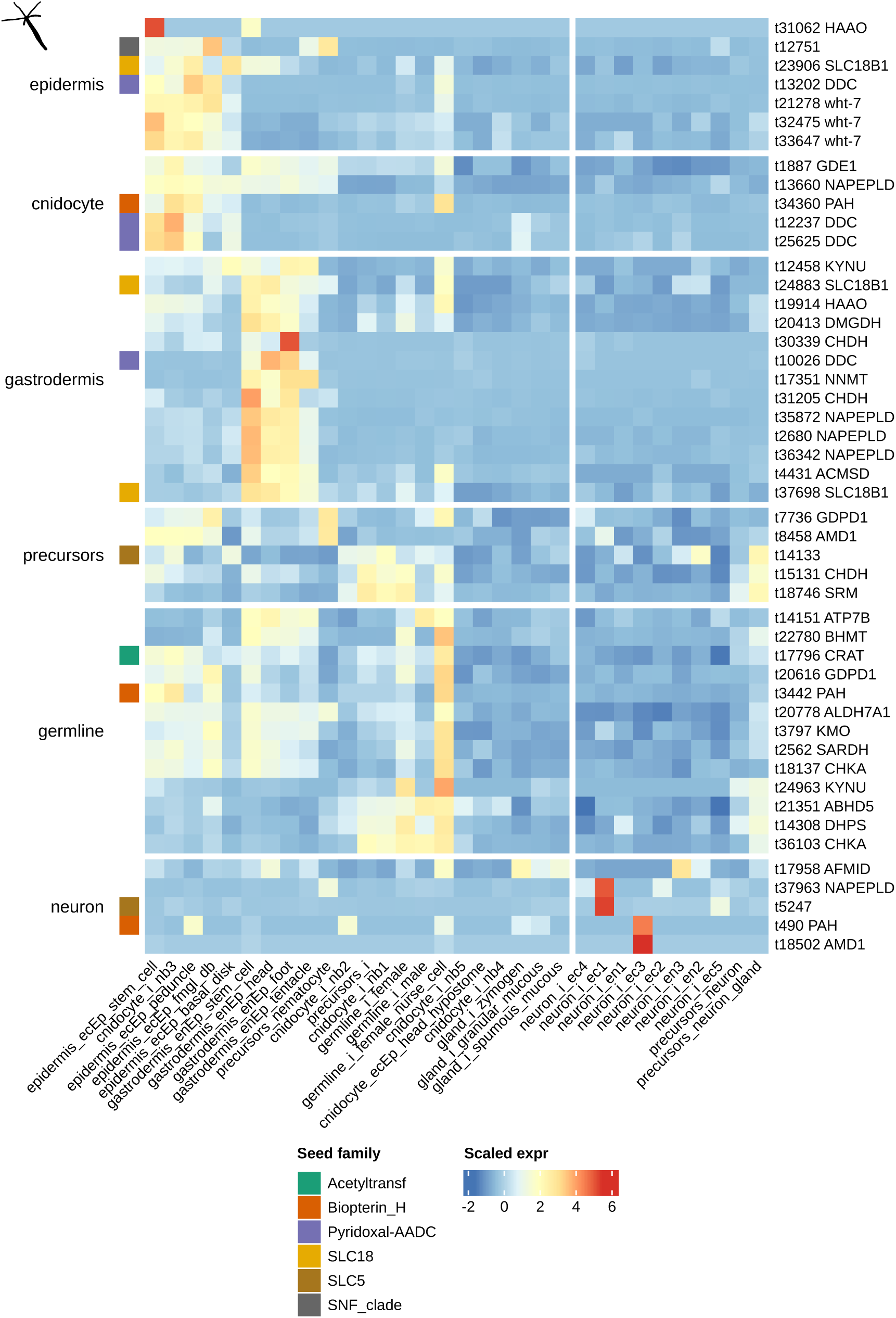
Monoamine metabolism­associated coregulon in *Hydra vulgaris*. Heatmap of the combined coregulon (genes co­expressed with the seed­family enzymes and transporters across cell types, intersected with GO:0009308 "amine metabolic process"; see Methods). Rows are genes (z­score­normalized across cell types), grouped by the cell­ type class in which their expression peaks (separated by white gaps) and clustered within each group; columns are cell types, ordered by hierarchical clustering of their transcriptomes (see Methods), with neuronal cell types on the right. Colored row annotation indicates the seed family of seed genes. The coregulons of the other neuron­bearing non­ bilaterians, *Nematostella vectensis* and *Mnemiopsis leidyi*, are shown in Fig. S11, and those of *Spongilla lacustris, Mus musculus* and *Drosophila melanogaster* in Figs. S12–S14.

The coregulons contain the expected core: the AAAH and AADC paralogs themselves, the choline-handling enzymes CHKA (choline kinase) and CHDH (choline dehydrogenase), CRAT (carnitine acetyltransferase), the polyamine vesicular transporter SLC18B1/VPAT, and DβH-related DOMON monooxygenases (MOXD1) in *Nematostella* but not in *Hydra*, consistent with the detection of octopamine in *Nematostella* only (Fig. 1; Discussion). All three species’ coregulons additionally contain orthologs of the ABC transporter white (ABCG family), which controls 5-HT, DA and histamine content in the *Drosophila* brain (*95*): three paralogs in *Hydra*, two in *Nematostella*, and one in *Mnemiopsis*. ATP7B, a copper transporter required for DβH function (*96, 97*), is co-regulated in *Hydra*.

Inspection of the coregulons highlighted three additional metabolic neighborhoods of potential interest. First, four kynurenine-pathway enzymes are present in the *Hydra* coregulon, namely KYNU and HAAO/3-HAO (two paralogs each), KMO and ACMSD, together with the carrier ATP7B and the lipid-handling NAPE-PLD/GDPD1/GDE1 module; CCBL2 (two paralogs) is present in *Mnemiopsis*, and TDO2 appears in *Mnemiopsis* and *Nematostella*. The *Spongilla* coregulon (Fig. S12) contains the most complete kynurenine pathway of any species in this study: IDO1/IDO2, two AFMID paralogs, KYNU, KMO, HAAO and CCBL2, covering every enzymatic step from tryptophan to 3-hydroxyanthranilate. The *Drosophila* coregulon (Fig. S14) additionally contains vermilion (TDO2), cinnabar (KMO) and kynurenine formamidase (AFMID), so the pathway is co-regulated with the amine-biosynthesis machinery in every invertebrate species we examined. This cross-species co-regulation suggests that tryptophan in these animals is partitioned between monoamine biosynthesis and kynurenine catabolism in a coordinated, cell-type-specific manner, with sponges and cnidarians retaining a more complete pre-bilaterian kynurenine repertoire than has previously been reported. Second, five NAPE-PLD paralogs together with two GDPD1 paralogs and GDE1, enzymes involved in N-acylethanolamine and anandamide biosynthesis, are present in the *Hydra* coregulon and NAPE-PLD is also present in *Mnemiopsis*, consistent with the previously reported endocannabinoid signaling in *Hydra* (*98*). Third, the *Nematostella* coregulon contains three paralogs of acetylserotonin O-methyltransferase (ASMT, also known as HIOMT), the terminal enzyme of melatonin biosynthesis, in line with earlier reports of melatonin in *Nematostella* (*99*), together with the ethanolamine-pathway enzyme Pect (phosphate cytidylyltransferase 2), the polyamine-biosynthesis enzyme deoxyhypusine synthase (DHPS) and the polyamine antizyme OAZ1; the *Hydra* coregulon additionally contains DHPS, the betaine-recycling enzyme BHMT, and the lipase regulator ABHD5; *Mnemiopsis* contains OAZ2. Together, these genes link the amine and choline pools to adjacent ethanolamine, polyamine and lipid metabolism, and point to additional small-molecule signaling and metabolic pathways co-deployed with the amine-biosynthesis machinery in non-bilaterian cells. Their functional significance remains to be tested.

## Discussion

Earlier analyses based on the bilaterian content of biosynthetic enzymes, transporters and receptors concluded that monoaminergic signaling is a bilaterian innovation (*40*). Our results argue otherwise, based on independent lines of evidence. Biogenic amines, including acetylcholine, are produced in sponges and cnidarians. The broad-specificity machinery responsible for their synthesis, transport and reception is present in these animals and predates animal multicellularity for some components. The dedicated, narrowly tuned bilaterian system is best understood as a specialization, through gene duplication, of an older promiscuous toolkit.

### Promiscuous biosynthesis explains the presence of monoamines in non-bilaterians

The metabolomic detection of tyramine, 2-phenylethylamine, tryptamine, dopamine, serotonin (*Hydra* only), histamine and adrenaline in *Spongilla lacustris, Hydra vulgaris* and *Nematostella vectensis* (Fig. 1) may be explained by the demonstrated promiscuity of the ancestral enzymes, despite the absence of dedicated bilaterian counterparts. AADC decarboxylates not only L-DOPA and 5-HTP but also tyrosine, phenylalanine, tryptophan and histidine, producing trace amines and histamine directly from their amino-acid precursors (*61–65, 100*). AADC orthologs are present in every non-bilaterian animal group we examined, and the *Trichoplax* ortholog can support tyramine synthesis in vivo (*42*). A single enzyme can thus provide a complete route to trace amines and histamine, although sponges also use decarboxylases outside this clade (*41*).

Serotonin and dopamine canonically require the dedicated hydroxylases TPH and TH, which are bilaterian-specific. The only biopterin-dependent hydroxylases present in non-bilaterians are PAH orthologs, whose primary substrate is phenylalanine. PAH from mouse (*59*), *Drosophila, Caenorhabditis* and several bacteria (*14, 60, 101*) also hydroxylates tryptophan to 5-HTP, indicating that broad-spectrum hydroxylase activity is an ancestral, not derived, property of this family. Consistent with this enzymatic promiscuity, PAH-dependent serotonin biosynthesis has been demonstrated in vivo in both mice and *C. elegans*, including serotonin production in the absence of canonical TPH activity (*59, 101*). The detection of serotonin in *Hydra* is therefore most parsimoniously explained by PAH supplying 5-HTP for AADC. Also, the occurrence of serotonin and serotonin-binding sites in cnidarians has been independently documented (*31, 36, 38, 102, 103*).

Dopamine is more challenging, because PAH hydroxylates tyrosine to L-DOPA only poorly (*104–107*). An alternative bilaterian route exists in which CYP2-family cytochrome P450 enzymes convert tyramine directly to dopamine (*93, 94*). CYP2 orthologs are present in non-bilaterian genomes (Fig. S1B), and we find CYP2D co-expressed with AADC in a candidate dopaminergic *Hydra* neuron type (Fig. 7B). L-DOPA was below the limit of quantification in all our samples, which fits the tyramine-to-dopamine route rather than the classical L-DOPA route, a prediction that can be tested directly by heterologous expression assays. For norepinephrine and octopamine, bilaterian DβH/TβH is replaced in non-bilaterians by clade-related MOXD1/MOXD2 co-orthologs (*108*), a likely source of the OCT we detect in *Nematostella* (not *Hydra*) and the NE inferred from epinephrine measurements. The single-cell coregulon of *Spongilla lacustris* reveals an even more pronounced co-regulation of this machinery: six co-expressed MOXD1-family paralogs (Fig. S12), the largest DβH-clade expansion in any species in this study. The retention of an expanded catecholamine β-monooxygenase repertoire in an animal that lacks neurons indicates that the enzymatic machinery for catecholamine production predates the nervous system, consistent with the metabolomic detection of dopamine and adrenaline in *Spongilla* (Fig. 1) and with dopamine responses and dopamine-related gene expression reported in other sponges (*109, 110*). Direct functional evidence now comes from *Spongilla lacustris*, where trace amines coordinate whole-body contractions through decarboxylases and vesicular transporters expressed in secretory cells, in the absence of neurons (*41*). Epinephrine itself is most plausibly produced via PNMT-like co-orthologs detectable in sponges and cnidarians (*111*).

In every case, the same evolutionary logic applies: a broadly active ancestral enzyme was complemented in Bilateria by a more specific duplicate that specialized in a reaction, while the ancestral enzyme remained with a multi-substrate functionality. Trace amines and histamine, which require only a single decarboxylation, were likely the earliest monoamines to acquire signaling roles. The two bilaterian reference coregulons in this study, of *Mus musculus* (Fig. S13) and *Drosophila melanogaster* (Fig. S14), display this logic directly: both contain dedicated, narrow-specificity genes (e.g., TPH1, NET and VMAT1 in mouse; Tβh, Tdc2, Trh, ChAT, VAChT, DAT and Vmat in *Drosophila*) that arose within the same families whose broad-specificity members (AADC, CrAT, PAH, MOXD1, SLC18B) we recover as the core amine-producing toolkit in sponges and cnidarians.

### Acetylcholine signaling is present in cnidarians

Acetylcholine is directly detected in *H. vulgaris* and *N. vectensis* (Fig. 1); muscarinic receptors orthologous to bilaterian mAChRs are present in medusozoan cnidarians (Fig. 4A); nicotinic receptor orthologs are present in both medusozoans and anthozoans (Fig. 5); and ACh immunoreactivity in *Nematostella* labels muscle-associated cells (Fig. 6). Two *Hydra* mAChR orthologs, ACh-rec1 and ACh-rec2, retain a fully conserved orthosteric site and are predicted to bind ACh as strongly as bilaterian mAChRs (Fig. 4B); their transcript is expressed predominantly in neurons (Fig. 4E).

The missing piece of a cnidarian cholinergic system has been the ACh biosynthetic enzyme: cnidarians lack ChAT (*112*). In mammals, extraneuronal ACh is produced not by ChAT but by CrAT, which acetylates choline as a side-activity of its primary reaction on carnitine (*44–47*), and a small number of substitutions in its substrate channel can flip the catalytic preference (*51, 55*). The *Hydra, Nematostella* and *Spongilla* CrATs retain residues associated with broad-specificity CrAT activity (Fig. 2B; Fig. S3), although some anemone paralogs likely exclude choline (Fig. S3D), and could therefore produce ACh as a minor by-product of carnitine metabolism. The ACh-to-acetylcarnitine ratios we measure are consistent with this: acetylcarnitine exceeds ACh in all three non-bilaterians, whereas the ratio is inverted in the bilaterian *Platynereis*, which has ChAT (Table S1). For transport and vesicular loading, the bilaterian-specific transporters VAChT, VMAT1 and VMAT2 are absent from cnidarians, but the broad-specificity transporters SLC22 (*67–69*) and SLC18B/VPAT (*71, 72*) are present and known to carry ACh and biogenic amines non-selectively.

### From by-product to specific signal

A coherent picture emerges. ACh is a thermodynamic by-product of CrAT activity wherever carnitine metabolism is active (*44–46, 52*): acetyl-CoA generated by fatty-acid oxidation equilibrates with acetylcarnitine via CrAT, and the same equilibrium reaction acting on choline rather than carnitine releases ACh. Neurons, with their elevated energy demand (*113*), are precisely such cells. The transporters that carry ACh into vesicles in non-bilaterians also carry monoamines, polyamines and carnitine itself (*72, 79, 114*); bilaterian VAChT and VMAT, which separate these pools, evolved by duplication from this shared precursor. The metabotropic receptors that recognize ACh in cnidarians are nested within a clade of monoamine receptors (Fig. 4A): mAChRs evolved by duplication within this aminergic clade, and the bilaterian specific dopamine, serotonin and trace-amine receptors evolved as further expansions within it. Independent support comes from Placozoa: *Trichoplax* orthologs of bilaterian melatonin receptors respond to non-acetylated trace amines rather than N-acetylated ligands (*42*), suggesting that the same logic, ancient promiscuous reception followed by bilaterian ligand specialization, applies across the whole aminergic GPCR clade. In pre-bilaterian animals, ACh and monoamines were therefore likely made opportunistically as metabolic by-products, co-released through shared transporters, and received by ligand-tolerant GPCRs, with biological specificity arising from the cellular context of reception rather than from dedicated presynaptic release.

### Specialization *in Bilateria*

The transition to bilaterian-style synaptic transmission is a series of duplications along the bilaterian stem: ChAT (choline-specific biosynthesis), VAChT/VMAT (separate vesicular pools), the high-affinity ChT (choline recycling; Fig. 3B) and AChE (rapid ACh removal). The embedding of VAChT within the ChAT locus established co-regulated expression of biosynthesis and vesicular loading (*115, 116*). Postsynaptic anchoring of nAChRs via Rapsyn (in muscle) and DLG4/PSD-95 (in central neurons) localized the signal, but the single-cell expression of muscle-type nAChR subunits in *Hydra* and *Nematostella* muscle cells (Figs. 5, S6, S7) shows that cholinergic neuromuscular signaling itself was already in place in the cnidarian–bilaterian ancestor and was specialized, not invented, in bilaterians. The secondary loss of cholinergic NMJs in *Drosophila*, where iGluRs replaced muscle nAChRs (Fig. S5), confirms the picture by counter-example. Biogenic-amine signaling is not a bilaterian innovation but a deep, pre-bilaterian mode of cellular communication that bilaterians inherited, separated and accelerated. Whether the dedicated monoaminergic neurons and synapses of bilaterians derive from these ancestral systems or recruited them independently remains open (*40, 117*).

## Methods

### Targeted Metabolomics by LC-MS/MS-MRM

For each replicate, 50 *Spongilla lacustris* gemmules (collected at Lake Constance, Germany) were hatched and grown in filtered lake water at 18 °C for 7 days in the dark. After a water exchange and an additional day of growth, the sponges were collected using a cell scraper. *Hydra vulgaris* (culture kindly provided by Prof. Dr. Suat Özbek, COS Heidelberg) were maintained in artificial *Hydra* medium (1 mM Tris-HCl, 1 mM CaCl2, 0.1 mM MgCl2, 0.1 mM KCl, 1 mM NaHCO3, pH 7.4) at 18 °C. Before extraction, ten animals were starved for 4 days for each replicate. *Nematostella vectensis* (animals kindly provided by Dr. Aissam Ikmi and Peter Steenbergen, Developmental Biology Unit, EMBL Heidelberg) were kept in 12 ppt artificial seawater (Sea Salt; Instant Ocean) at 16 °C. Before collection, the animals were starved for 4 days. For each replicate, one adult specimen was collected. *Platynereis dumerilii* were cultured in natural sea water under a constant temperature of 16 °C and a 16:8 h light:dark cycle. One atoke worm (adult, sexually immature) of about 3 cm length was collected for each replicate.

All samples were collected in 2 mL cryotubes (Biosphere plus; Sarstedt, Germany) and immediately flash frozen in liquid nitrogen. Additionally, the respective surrounding culture media were collected as controls. Samples were homogenized via bead beating (FastPrep-24 Classic (MP Biomedicals; USA), 5 x 20 s at 6 m/s) using 2.3 mm zirconia beads (Carl Roth, Germany) in extraction solution (250 µL methanol, 2% formic acid, (Sigma-Aldrich, Germany)). Homogenates were centrifuged (Centrifuge 5145 R; Eppendorf, Germany) for 15 min at 16,100 rcf and 4 °C. Afterwards, supernatants were transferred to silanized glass microinserts (200 µL) in 1.5 mL analytical vials, which were closed with polypropylene caps. The samples were placed into the autosampler (maintained at 4 °C) until the analysis, which was conducted on the same day.

### LC-MS/MS parameters

The analysis was performed on an ExionLC AD UHPLC system coupled to a QTRAP 6500+ triple quadrupole mass spectrometer (Sciex, MA, USA) in positive ESI (electrospray ionization) mode. Chromatographic separation was carried out on an Atlantis Premier BEH Z-HILIC column (2.1 mm × 100 mm, 1.7 µm; Waters, MA, USA) at a flow rate of 0.3 mL/min. The mobile phase consisted of water (mobile phase A) and acetonitrile:water (9:1; mobile phase B), which were modified with a total buffer concentration of 10 mM ammonium formate (Sigma-Aldrich, Germany), respectively. The aqueous portion of each mobile phase was adjusted to pH 3.0 via addition of formic acid. The following gradient (13 min total run time including re-equilibration) was applied (min/%B): 0/100, 2/100, 8/80, 9/60, 10/60, 10.5/100, 13/100. Column temperature was maintained at 40 °C and sample injection volume was set to 5 µL. Ion source parameters were set to the following values: ion spray voltage: 5000 V, nebulizer gas (zero-grade air): 80 psi, heater gas (zero-grade air): 40 psi, curtain gas (nitrogen): 0 psi, heater temperature: 350 °C.

Detection of target analytes was conducted using manually tuned precursor-fragment mass transition in scheduled multiple reaction monitoring (sMRM) mode (Table S6). Absolute quantification of the analytes was achieved by standard addition in true matrix. Three defined concentration levels of each metabolite were obtained by spiking low volumes to 25 µL aliquots of each replicate. Absolute concentration values were calculated via the resulting standard addition lines (derived from integrated peak areas in SciexOS software). Standard addition is independent of potential matrix effects and allowed the affirmation of metabolite identification in the samples, which was also verified via injection of reference standards in neat solution. Concentrations reported for phenylethylamine, tyramine and tryptamine in the case of *Nematostella vectensis* represent minimal values. In these cases, the sample concentrations measured were outside of the linear range of the standard addition.

### Immunohistochemistry of Acetylcholine in *Nematostella vectensis*

Adult polyps were relaxed in 7% w/v MgCl2 for about 10 minutes and then cut open longitudinally with scissors to obtain pieces of the body wall spanning at least two parietal muscle tracts. Animals were fixed for 120 minutes at 4 °C in 4% paraformaldehyde (PFA) in *Nematostella* medium. After incubation animals were washed five times in PBS containing 0.2% Triton X-100 (PTx). Samples and primary antibodies were blocked in 80% PTx, 20% sheep serum, 1% BSA for 2 hours on a shaker. The blocking solution was then replaced by the antibody solution and samples were incubated at 4 °C over the weekend. Primary antibodies against ACh (Merck/Millipore, AB5522) were used at a dilution of 1:250. After incubation, samples were washed five times in PTx for at least 5–10 minutes. Samples were then incubated overnight at 4 °C in fluorophore-conjugated secondary antibody (Invitrogen, A11011) diluted 1:500 in blocking solution. Samples were then washed three times in PTx and twice in PBS. Samples were either incubated in 3 µl phalloidin/100 µl PBS and 0.75 µl DAPI/1 ml PBS overnight or in 0.75 µl DAPI/1 ml PBS for 2 h. Body wall pieces were gradually infiltrated in 30%, 50%, 86% glycerol and left in 86% overnight and then embedded in Fluoromount™ Aqueous Mounting Medium from Sigma-Aldrich, to prevent loss of fluorescence. Micrographs were taken at a Nikon Deconvolution microscope or at the Leica SP5 and post-processed with Adobe Photoshop.

### Structural binding-affinity prediction with Boltz-2

To evaluate the predicted ligand specificity of the *H. vulgaris* mAChR orthologs and of representative bilaterian and non-bilaterian receptors, we used Boltz-2 (*86*) to compute structural binding-affinity predictions for each receptor against six small-molecule ligands: acetylcholine, histamine, dopamine, serotonin, tryptamine and 2-phenylethylamine. Receptor sequences were submitted to Boltz-2 (v2.2.1) in default mode, with multiple sequence alignments generated by the ColabFold MMseqs2 server, and per-receptor-per-ligand predictions were obtained as JSON outputs containing two key metrics: a continuous binding affinity score (affinity_pred_value, lower = stronger binding) and a binary binding probability (affinity_probability_binary, 0–1). The reported values are the mean of the two prediction heads; the values of both heads are given in Table S4. As positive controls we used the human, mouse and *Drosophila* muscarinic receptors with acetylcholine, the human and mouse histamine receptors HRH1–4 with histamine, and the human serotonin (5-HT2A–C) and dopamine (DRD2–4) receptors with their cognate ligands; as negative controls we used MneR (*Mnemiopsis leidyi*) and TriR (*Trichoplax adhaerens*) against all six ligands (Table S4). TriR is distinct from the monoamine-responsive placozoan receptors characterized in (*42*).

### Sequence Datasets

We assembled a protein sequence collection of 69 genomic and transcriptomic datasets of 68 eukaryotic species, with an emphasis on the comprehensive taxonomic representation of all major metazoan clades, unicellular animal relatives and fungal species, complemented by representatives of various major groups of eukaryotes (Table S2). For those cases with a published genome available in major sequence databases (e.g. Uniprot Proteomes, Ensembl), we used the latest peptide prediction after filtering, when possible, for the longest isoform per gene (Uniprot provides directly the longest isoform peptide set). Given that many taxonomically important clades are vastly undersampled and no genome is available, we also used published transcriptomic data, or yet unpublished data kindly provided by collaborators within the EvoCell network. Particularly for those species with available single-cell expression data, special attention was given to matching the species’ transcriptome assembly and annotation version to those used in the single-cell analysis. For *Hydra vulgaris* only, both a genome to use as reference, and a transcriptome that matches the single-cell data, were used. In parallel, we downloaded from Uniprot the prokaryotic Reference Proteomes (representatives of each group).

### Phylogenetic reconstruction analysis methods

For all the protein families analyzed in this work, we used a base approach which was further adapted to each case’s special characteristics after an initial evaluation. We screened the literature for all the known genes involved in the biosynthesis, vesicular transport and reuptake, and reception of neurotransmitters, and scaffolding proteins important for the postsynaptic anchoring of receptors. For each of these genes we found the family which they belong to, based on the Pfam annotation of their protein products. In the cases where the proteins were annotated with more than one Pfam domain, we used the most specific one of the family we were interested in. The hmm profile of each Pfam domain family was used to screen our eukaryotic species database, as well as the Uniprot prokaryotic Reference Proteomes for proteins containing the domain. The hmmsearch (HMMER 3.3.2 (*118*)) with the--cut_ga (gathering cutoff) parameter was used for all searches. For all the cases with detected prokaryotic homologs, we used a taxonomy-aware strategy to deal with redundancy and subsample the prokaryotic sequences to include in the subsequent steps. If the number of prokaryotic hits exceeded 300, we selected a maximum of 100 sequences per bacterial or archaeal phylum. For the two vast multi-member superfamilies, the MFS and GPCR-A, we used the following similarity-based clustering approach: first an all-vs-all blastp search (Protein-Protein BLAST 2.11.0+) was performed using an E-value threshold of 1e-5. The negative base 10 logarithm of each query-target E-value was fed into the MCL (*119*) clustering software, which breaks down the big network of sequence similarity across the family into clusters of closely related sequences. Then, we extracted those clusters which contained the proteins (families) of interest. In both cases we used different inflation parameters (controlling clustering granularity) and selected the optimal number for each superfamily (1.4 for MFS, 6.0 for GPCR-A). Our maximum likelihood phylogenetic reconstruction pipeline was decided on a case-by-case basis, depending on the size of each protein set to be processed, as well as after a first evaluation of the initial results. The standard pipeline that we used included MAFFT (v7.475) (*120*) with an iterative refinement method using WSP and consistency scores (E-INS-i or L-INS-i) for multiple sequence alignment, alignment columns that contained more than 90% gaps were removed using a custom script, and a maximum likelihood phylogenetic tree was computed with IQ-TREE (v1.6.12) (*121*) with model selection out of all available LG models, and ultrafast bootstrap approximation (*122*) for computing support values. ETE toolkit (*123*) was used in all cases for tree processing, taxonomic annotation, and tree visualization for inspection. Alignment methods were selected case by case after initial testing; besides MAFFT, we used Clustal Omega (v1.2.4) (*124*) with default parameters, and for some very large sets MAFFT with default parameters. For some large datasets we removed columns with more than 99% gaps, as e.g. a uniquely conserved region in less than 100 sequences in an alignment of 1,000 sequences in total would be removed with the 90% threshold. In such cases, an ML tree was computed and evaluated using FastTree (v2.1.11 SSE3; all runs with the LG model, option -lg) (*125*). In the presence of multiple families forming distinct monophyletic clades in the phylogenetic tree, the sequences within the monophyletic clade of interest were extracted, and another tree with the standard more accurate algorithm was computed; for some large clades (SLC22, CYP2 and the cationic Cys-loop subunits) this tree was computed on a reduced, representative taxon sampling (Table S5). The alignments and trees underlying each figure, with the number of sequences, alignment method, gap filtering, tree-inference method and substitution model, are listed in Table S5.

### Single-cell expression Datasets and analysis

Quality filtered UMI tables and authors-derived cell type assignments and annotations were collected for whole body single-cell datasets for mouse (*87*), fly (no head) (*126*), the cnidarians *Hydra vulgaris* (*88*) and *Nematostella vectensis* (*127*), the ctenophore *Mnemiopsis leidyi* (*128*) and the sponge *Spongilla lacustris* (*129*). For *Hydra* and *Nematostella* we used the coarse and fine cell-type annotations of Levy et al. (*130*). UMI tables were averaged per cell type prior to all downstream analyses. In all heatmaps, cell types were ordered by complete-linkage hierarchical clustering of their averaged transcriptomes (distance 1 − Pearson r), with neuronal cell types placed on the right.

#### Co-expression analysis of amine-metabolism genes

For each species, the normalized cell-type-averaged expression matrix produced above was used to compute pairwise Pearson correlations between all genes (using rcorr from the Hmisc R package; missing p-values were set to zero). For each seed gene, defined as a member of the eight protein families relevant to amine biosynthesis, transport and reuptake covered in this study (carnitine/choline acetyltransferases; biopterin-dependent hydroxylases; pyridoxal-AADC decarboxylases; DOMON-domain monooxygenases; cytochrome P450-CYP2D6 clade; SLC18, SLC5 and SNF transporter clades), we extracted the set of genes whose Pearson r with the seed was ≥ 0.5 and whose Benjamini–Hochberg-adjusted two-tailed p-value was ≤ 0.05. The BH correction was applied within the per-species candidate pool (GO:0009308 genes plus the seed-family members), the relevant hypothesis space for an amine coregulon, and seeds were retained only if they had at least one non-self correlated partner at this threshold. The resulting per-seed gene lists were intersected with the GO:0009308 ("amine metabolic process") annotation, transferred from eggNOG-mapper-derived orthology assignments. The per-seed amine-GO gene lists were then merged, together with the retained seeds, into a single per-species set (the "coregulon" of the manuscript), which was visualized as a heatmap (ComplexHeatmap; row z-scores) with genes grouped by the cell-type class in which their expression peaks and clustered within each group (complete linkage, distance 1 − Pearson r). For brevity we use the term coregulon throughout for the co-expressed gene set; we note that the analysis measures co-expression patterns across cell-type-averaged profiles rather than coregulation in the strict gene-regulatory sense.

#### Structural prediction, visualization and alignment

Structures for *Nematostella, Hydra* and *Spongilla* CrATs were predicted using ColabFold in default mode (*57*). Protein structures were visualized in PyMOL (v2.3.5; Schrodinger, Inc.). Protein superpositions were created using the *super* command. Electrostatic surface properties of the proteins in aqueous solutions were calculated using the APBS Electrostatics plugin (*131*). Structure alignment (Table S3) of the ColabFold predictions and protein crystal structures to the AlphaFold predictions of SwissProt was done by Foldseek (*132*) using default parameters. Residues discriminating between protein subfamilies (ChAT versus the other carnitine acyltransferases, Fig. S3C; the cnidarian–bilaterian mAChR clade versus the other GPCRs, Fig. 4C) were identified with the Sequence Harmony method (*56*) as implemented in Jalview (*133*).

## Supporting information

Table S1 targeted metabolomics

Table S2 species sequence datasets

Table S3 Foldseek search results

Table S4 Boltz-2 binding predictions

Table S5 alignments and phylogenetic trees

Table S6 targeted metabolomics MRM parameters

## Data and code availability

Multiple sequence alignments and phylogenetic trees (Table S5), eggNOG-mapper annotations, cell-type-averaged single-cell expression matrices, coregulon tables, and Boltz-2 inputs and predictions (Table S4) are available at Zenodo (DOI: 10.5281/zenodo.22813072). Code to reproduce the analyses and figures is available at https://github.com/alxndrsPittis/neurotransevo (frozen for this submission at tag v1.0.0; archived at Zenodo, DOI: 10.5281/zenodo.23324945).

## Acknowledgements

We thank Prof. Dr. Suat Özbek (Centre for Organismal Studies, Heidelberg) for kindly providing a *Hydra vulgaris* culture.

We thank Dr. Aissam Ikmi and Peter Steenbergen (Developmental Biology Unit, EMBL Heidelberg) for kindly providing *Nematostella* adults.

We thank Dr. Bernhard Drotleff (Metabolomics Core Facility, EMBL Heidelberg) for the thorough and quick targeted metabolomic experiments.

We thank the EvoCell community for kindly providing unpublished sequence data.

## Author contributions

D.A., J.M.M., A.A.P. and J.H.C. conceived the study. A.A.P. designed and performed all computational and phylogenetic analyses. F.R. performed the structural modeling and the targeted metabolomics experiments. A.G.C. and U.T. performed the immunostaining experiments. L.A.Y.-G. and G.J. contributed to the conceptual framework and the interpretation of the results. D.T. and M.M. contributed to the experimental work and discussions. All authors contributed to the writing and approved the final manuscript.

## References

1. O.-B. Tysnes, A. Storstein, Epidemiology of Parkinson’s disease. J. Neural Transm. 124, 901–905 (2017).

2. S. A. Azizi, Monoamines: Dopamine, Norepinephrine, and Serotonin, Beyond Modulation, “Switches” That Alter the State of Target Networks. Neuroscientist 28, 121– 143 (2022).

3. A. Y. Deutch, “Neurotransmitters” in Fundamental Neuroscience (Elsevier, 2013), pp. 117–138.

4. R. Dalangin, A. Kim, R. E. Campbell, The Role of Amino Acids in Neurotransmission and Fluorescent Tools for Their Detection. Int. J. Mol. Sci. 21, 6197 (2020).

5. G. Pándy-Szekeres, J. Caroli, A. Mamyrbekov, A. A. Kermani, G. M. Keserű, A. J. Kooistra, D. E. Gloriam, GPCRdb in 2023: state-specific structure models using AlphaFold2 and new ligand resources. Nucleic Acids Res. 51, D395–D402 (2023).

6. R. Fredriksson, M. C. Lagerström, L.-G. Lundin, H. B. Schiöth, The G-protein-coupled receptors in the human genome form five main families. Phylogenetic analysis, paralogon groups, and fingerprints. Mol Pharmacol 63, 1256–1272 (2003).

7. M. Selcho, D. Pauls, B. El Jundi, R. F. Stocker, A. S. Thum, The role of octopamine and tyramine in Drosophila larval locomotion. J. Comp. Neurol. 520, 3764–3785 (2012).

8. P. Ji, F. Xu, B. Huang, Y. Li, L. Li, G. Zhang, Molecular Characterization and Functional Analysis of a Putative Octopamine/Tyramine Receptor during the Developmental Stages of the Pacific Oyster, Crassostrea gigas. PLoS One 11, e0168574 (2016).

9. S. Gonçalves, D. Nunes-Costa, S. M. Cardoso, N. Empadinhas, J. D. Marugg, Enzyme Promiscuity in Serotonin Biosynthesis, From Bacteria to Plants and Humans. Front. Microbiol. 13, 873555 (2022).

10. R. Marchiosi, A. R. Soares, J. Abrahão, W. D. dos Santos, O. Ferrarese-Filho, “L-DOPA and Dopamine in Plant Metabolism” in Neurotransmitters in Plant Signaling and Communication (Springer International Publishing, Cham, 2020), pp. 141–167.

11. Q. Liu, T. Gao, W. Liu, Y. Liu, Y. Zhao, Y. Liu, W. Li, K. Ding, F. Ma, C. Li, Functions of dopamine in plants: a review. Plant Signal. Behav. 15, 1827782 (2020).

12. E. C. Azmitia, “Evolution of serotonin: Sunlight to suicide” in Handbook of Behavioral Neuroscience (Elsevier, 2010), pp. 3–22.

13. I. Wessler, H. Kilbinger, F. Bittinger, C. J. Kirkpatrick, The biological role of non-neuronal acetylcholine in plants and humans. Jpn. J. Pharmacol. 85, 2–10 (2001).

14. J. Jiao, Y. Xia, Y. Zhang, X. Wu, C. Liu, J. Feng, X. Zheng, S. Song, T. Bai, C. Song, M. Wang, H. Pang, Phenylalanine 4-Hydroxylase Contributes to Endophytic Bacterium Pseudomonas fluorescens’ Melatonin Biosynthesis. Front. Genet. 12, 746392 (2021).

15. J. Cao, F. Shi, X. Liu, G. Huang, M. Zhou, Phylogenetic analysis and evolution of aromatic amino acid hydroxylase. FEBS Lett. 584, 4775–4782 (2010).

16. E. Hedner, M. Sjögren, P. A. Frändberg, T. Johansson, U. Göransson, M. Dahlström, P. Jonsson, F. Nyberg, L. Bohlin, Brominated cyclodipeptides from the marine sponge Geodia barretti as selective 5-HT ligands. J. Nat. Prod. 69, 1421–1424 (2006).

17. K. Ellwanger, M. Nickel, Neuroactive substances specifically modulate rhythmic body contractions in the nerveless metazoon Tethya wilhelma (Demospongiae, Porifera). Front. Zool. 3, 7 (2006).

18. A. M. Sokolova, E. E. Voronezhskaya, Dopamine-like immunoreactivity in sponge larvae. Invertebr. Zool. 18, 345–354 (2021).

19. M. Anctil, Cholinergic and monoaminergic mechanisms associated with control of bioluminescence in the ctenophore *Mnemiopsis leidyi*. J. Exp. Biol. 119, 225–238 (1985).

20. M. Jager, C. Dayraud, A. Mialot, E. Quéinnec, H. le Guyader, M. Manuel, Evidence for involvement of Wnt signalling in body polarities, cell proliferation, and the neuro-sensory system in an adult ctenophore. PLoS One 8, e84363 (2013).

21. L. L. Moroz, K. M. Kocot, M. R. Citarella, S. Dosung, T. P. Norekian, I. S. Povolotskaya, A. P. Grigorenko, C. Dailey, E. Berezikov, K. M. Buckley, A. Ptitsyn, D. Reshetov, K. Mukherjee, T. P. Moroz, Y. Bobkova, F. Yu, V. V. Kapitonov, J. Jurka, Y. V. Bobkov, J. J. Swore, D. O. Girardo, A. Fodor, F. Gusev, R. Sanford, R. Bruders, E. Kittler, C. E. Mills, J. P. Rast, R. Derelle, V. V. Solovyev, F. A. Kondrashov, B. J. Swalla, J. V. Sweedler, E. I. Rogaev, K. M. Halanych, A. B. Kohn, The ctenophore genome and the evolutionary origins of neural systems. Nature 510, 109–114 (2014).

22. M. Carlberg, Localization and identification of catechol compounds in the ctenophore Mnemiopsis leidyi. Comp. Biochem. Physiol. C 91, 69–74 (1988).

23. K. Hanai, M.-A. Kitajima, Two types of surface amine receptors modulating the feeding response in Hydra japonica: the depressing action of dopamine and related amines. Chem. Senses 9, 355–367 (1984).

24. D. Hudman, I. D. McFarlane, The role of L-Dopa in the nervous system of sea anemones: a putative inhibitory transmitter in tentacles. J. Exp. Biol. 198, 1045–1050 (1995).

25. N. Isomura, C. Yamauchi, Y. Takeuchi, A. Takemura, Does dopamine block the spawning of the acroporid coral Acropora tenuis? Sci. Rep. 3, 2649 (2013).

26. J. A. Westfall, S. R. Elliott, P. S. MohanKumar, R. W. Carlin, Immunocytochemical evidence for biogenic amines and immunogold labeling of serotonergic synapses in tentacles of Aiptasia pallida (Cnidaria, Anthozoa). Invertebr. Biol. 119, 370–378 (2000).

27. H. Q. Marlow, M. Srivastava, D. Q. Matus, D. Rokhsar, M. Q. Martindale, Anatomy and development of the nervous system of Nematostella vectensis, an anthozoan cnidarian. Dev. Neurobiol. 69, 235–254 (2009).

28. M. Carlberg, N. Mons, M. Geffard, D. R. Nässel, L-DOPA and FMRFamide immunoreactivity in the tentacular nerve plexus of the sea anemone Metridium senile. Comp. Biochem. Physiol. C 94, 435–440 (1989).

29. M. Carlberg, R. Elofsson, Presence of 3,4-dihydroxyphenylalanine and 3,4,5-trihydroxyphenylalanine in a coelenterate nervous system: Possible tyrosinase-mediated accumulation. Neurochem. Int. 11, 161–167 (1987).

30. J. Taira, I. Higa, E. Tsuchida, N. Isomura, A. Iguchi, Neurotransmitters in hermatypic coral, Acropora spp., and its contribution to synchronous spawning during reproductive event. Biochem. Biophys. Res. Commun. 501, 80–84 (2018).

31. M. Carlberg, E. Rosengren, Biochemical basis for adrenergic neurotransmission in coelenterates. J. Comp. Physiol. B 155, 251–255 (1985).

32. E. W. Awad, M. Anctil, Identification of β-like adrenoceptors associated with bioluminescence in the sea pansy Renilla koellikeri. J. Exp. Biol. 177, 181–200 (1993).

33. M. Anctil, P. Hurtubise, M.-A. Gillis, Tyrosine hydroxylase and dopamine-beta-hydroxylase immunoreactivities in the cnidarian Renilla koellikeri. Cell Tissue Res. 310, 109–117 (2002).

34. D. Umbriaco, M. Anctil, L. Descarries, Serotonin-immunoreactive neurons in the cnidarian Renilla koellikeri. J. Comp. Neurol. 291, 167–178 (1990).

35. A. K. Pani, M. Anctil, D. Umbriaco, Neuronal localization and evoked release of norepinephrine in the cnidarian Renilla koellikeri. J. Exp. Zool. 272, 1–12 (1995).

36. A. K. Pani, M. Anctil, Evidence for biosynthesis and catabolism of monoamines in the sea pansy Renilla koellikeri (Cnidaria). Neurochem. Int. 25, 465–474 (1994).

37. G. Zega, R. Pennati, A. Fanzago, F. De Bernardi, Serotonin involvement in the metamorphosis of the hydroid Eudendrium racemosum. Int. J. Dev. Biol. 51, 307–313 (2007).

38. D. W. McCauley, Serotonin plays an early role in the metamorphosis of the hydrozoan Phialidium gregarium. Dev. Biol. 190, 229–240 (1997).

39. N. Takeda, C. N. Svendsen, Monoamine concentrations in Hydra magnipapillata. Hydrobiologia 216-217, 549–554 (1991).

40. M. Goulty, G. Botton-Amiot, E. Rosato, S. G. Sprecher, R. Feuda, The monoaminergic system is a bilaterian innovation. Nat. Commun. 14, 3284 (2023).

41. R. X. Zang, N. Malaiwong, L. Wang, J. D. Maziarz, K. Jia, B. Drotleff, F. Stein, M. Noor, C. J. Roberts, M. Rettel, J. J. Schwarz, A. Ikmi, S. Watanabe, R. Prevedel, M. P. O’Donnell, J. M. Musser, An ancient monoaminergic signaling system coordinates contractility in a nerveless sponge, bioRxiv (2026). 10.64898/2026.02.14.704838.

42. L. A. Yañez-Guerra, T. D. Mayorova, J. Duruz, I. Muralidhara, I. Hardege, L. Guan, Y. Hongo, H. Nakano, H. Wijnen, A. Senatore, H. Watanabe, G. Jékely, Prebilaterian origin of monoaminergic signaling. Curr. Biol. 36, 3719–3733.e8 (2026).

43. R. R. Ramsay, J. H. Naismith, A snapshot of carnitine acetyltransferase. Trends Biochem. Sci. 28, 343–346 (2003).

44. S. Tuček, Regulation of acetylcholine synthesis in the brain. J. Neurochem. 44, 11–24 (1985).

45. J. Slavíková, S. Tuček, Choline acetyltransferase in the heart of adult rats. Pflugers Arch. 392, 225–229 (1982).

46. H. L. White, J. C. Wu, Choline and carnitine acetyltransferases of heart. Biochemistry 12, 841–846 (1973).

47. R. Roskoski Jr, H. E. Mayer, P. G. Schmid, Choline acetyltransferase activity in guinea-pig heart in vitro. J. Neurochem. 23, 1197–1200 (1974).

48. R. Berman, I. B. Wilson, D. Nachmansohn, Choline acetylase specificity in relation to biological function. Biochim. Biophys. Acta 12, 315–324 (1953).

49. J. Schuberth, Choline acetyltransferase purification and effect of salts on the mechanism of the enzyme-catalyzed reaction. Biochim. Biophys. Acta 122, 470–481 (1966).

50. A. G. Cordente, E. López-Viñas, M. I. Vázquez, J. H. Swiegers, I. S. Pretorius, P. Gómez-Puertas, F. G. Hegardt, G. Asins, D. Serra, Redesign of carnitine acetyltransferase specificity by protein engineering. J. Biol. Chem. 279, 33899–33908 (2004).

51. A. G. Cordente, E. López-Viñas, M. I. Vázquez, P. Gómez-Puertas, G. Asins, D. Serra, F. G. Hegardt, Mutagenesis of specific amino acids converts carnitine acetyltransferase into carnitine palmitoyltransferase. Biochemistry 45, 6133–6141 (2006).

52. A.-R. Kim, R. J. Rylett, B. H. Shilton, Substrate binding and catalytic mechanism of human choline acetyltransferase. Biochemistry 45, 14621–14631 (2006).

53. G. Jogl, Y.-S. Hsiao, L. Tong, Structure and function of carnitine acyltransferases. Ann. N. Y. Acad. Sci. 1033, 17–29 (2004).

54. K. D. Green, V. R. Porter, Y. Zhang, S. Garneau-Tsodikova, Redesign of cosubstrate specificity and identification of important residues for substrate binding to hChAT. Biochemistry 49, 6219–6227 (2010).

55. C. N. Cronin, Redesign of choline acetyltransferase specificity by protein engineering. J. Biol. Chem. 273, 24465–24469 (1998).

56. B. W. Brandt, K. A. Feenstra, J. Heringa, Multi-Harmony: detecting functional specificity from sequence alignment. Nucleic Acids Res. 38, W35–40 (2010).

57. M. Mirdita, K. Schütze, Y. Moriwaki, L. Heo, S. Ovchinnikov, M. Steinegger, ColabFold: making protein folding accessible to all. Nat. Methods 19, 679–682 (2022).

58. J. Jumper, R. Evans, A. Pritzel, T. Green, M. Figurnov, O. Ronneberger, K. Tunyasuvunakool, R. Bates, A. Žídek, A. Potapenko, A. Bridgland, C. Meyer, S. A. A. Kohl, A. J. Ballard, A. Cowie, B. Romera-Paredes, S. Nikolov, R. Jain, J. Adler, T. Back, S. Petersen, D. Reiman, E. Clancy, M. Zielinski, M. Steinegger, M. Pacholska, T. Berghammer, S. Bodenstein, D. Silver, O. Vinyals, A. W. Senior, K. Kavukcuoglu, P. Kohli, D. Hassabis, Highly accurate protein structure prediction with AlphaFold. Nature 596, 583–589 (2021).

59. A. Mordhorst, P. Dhandapani, S. Matthes, V. Mosienko, M. Rothe, M. Todiras, J. Self, W.-H. Schunck, A. Schütz, M. Bader, N. Alenina, Phenylalanine hydroxylase contributes to serotonin synthesis in mice. FASEB J. 35, e21648 (2021).

60. S. Alcañiz, F. J. Silva, Phenylalanine hydroxylase participation in the synthesis of serotonin and pteridines in Drosophila melanogaster. Comp. Biochem. Physiol. C 116, 205–212 (1997).

61. S. Udenfriend, C. T. Clark, E. Titus, 5-hydroxytryptophan decarboxylase: A new route of metabolism of tryptophan. J. Am. Chem. Soc. 75, 501–502 (1953).

62. W. Lovenberg, H. Weissbach, S. Udenfriend, Aromatic L-amino acid decarboxylase. J. Biol. Chem. 237, 89–93 (1962).

63. E. Werle, D. Aures, [On the purification and specificity of DOPA-decarboxylase]. Hoppe Seylers Z. Physiol. Chem. 316, 45–60 (1959).

64. J. G. Christenson, W. Dairman, S. Udenfriend, Preparation and properties of a homogeneous aromatic L-amino acid decarboxylase from hog kidney. Arch. Biochem. Biophys. 141, 356–367 (1970).

65. J. M. Saavedra, J. Axelrod, Effect of drugs on the tryptamine content of rat tissues. J. Pharmacol. Exp. Ther. 185, 523–529 (1973).

66. D. Drew, R. A. North, K. Nagarathinam, M. Tanabe, Structures and General Transport Mechanisms by the Major Facilitator Superfamily (MFS). Chem. Rev. 121, 5289–5335 (2021).

67. S. K. Nigam, The SLC22 Transporter Family: A Paradigm for the Impact of Drug Transporters on Metabolic Pathways, Signaling, and Disease. Annu. Rev. Pharmacol. Toxicol. 58, 663–687 (2018).

68. C. Zhu, K. B. Nigam, R. C. Date, K. T. Bush, S. A. Springer, M. H. Saier Jr, W. Wu, S. K. Nigam, Evolutionary Analysis and Classification of OATs, OCTs, OCTNs, and Other SLC22 Transporters: Structure-Function Implications and Analysis of Sequence Motifs. PLoS One 10, e0140569 (2015).

69. D. C. Engelhart, J. C. Granados, D. Shi, M. H. Saier Jr, M. E. Baker, R. Abagyan, S. K. Nigam, Systems Biology Analysis Reveals Eight SLC22 Transporter Subgroups, Including OATs, OCTs, and OCTNs. Int. J. Mol. Sci. 21, 1791 (2020).

70. T. Nakata, T. Matsui, K. Kobayashi, Y. Kobayashi, N. Anzai, Organic cation transporter 2 (SLC22A2), a low-affinity and high-capacity choline transporter, is preferentially enriched on synaptic vesicles in cholinergic neurons. Neuroscience 252, 212–221 (2013).

71. M. Hiasa, T. Miyaji, Y. Haruna, T. Takeuchi, Y. Harada, S. Moriyama, A. Yamamoto, H. Omote, Y. Moriyama, Identification of a mammalian vesicular polyamine transporter. Sci. Rep. 4, 1–9 (2014).

72. Y. Moriyama, R. Hatano, S. Moriyama, S. Uehara, Vesicular polyamine transporter as a novel player in amine-mediated chemical transmission. Biochim. Biophys. Acta Biomembr. 1862, 183208 (2020).

73. Y. Kido, P. Matsson, K. M. Giacomini, Profiling of a prescription drug library for potential renal drug-drug interactions mediated by the organic cation transporter 2. J. Med. Chem. 54, 4548–4558 (2011).

74. D. Kobayashi, I. Tamai, Y. Sai, K. Yoshida, T. Wakayama, Y. Kido, J.-I. Nezu, S. Iseki, A. Tsuji, Transport of carnitine and acetylcarnitine by carnitine/organic cation transporter (OCTN) 2 and OCTN3 into epididymal spermatozoa. Reproduction 134, 651– 658 (2007).

75. Y. Kobayashi, N. Ohshiro, R. Sakai, M. Ohbayashi, N. Kohyama, T. Yamamoto, Transport mechanism and substrate specificity of human organic anion transporter 2 (hOat2 [SLC22A7]). J. Pharm. Pharmacol. 57, 573–578 (2005).

76. H. Koepsell, K. Lips, C. Volk, Polyspecific organic cation transporters: structure, function, physiological roles, and biopharmaceutical implications. Pharm. Res. 24, 1227– 1251 (2007).

77. H. Koepsell, Substrate recognition and translocation by polyspecific organic cation transporters. Biol. Chem. 392, 95–101 (2011).

78. D. C. Li, C. G. Nichols, M. Sala-Rabanal, Role of a Hydrophobic Pocket in Polyamine Interactions with the Polyspecific Organic Cation Transporter OCT3. J. Biol. Chem. 290, 27633–27643 (2015).

79. H. Koepsell, The SLC22 family with transporters of organic cations, anions and zwitterions. Mol. Aspects Med. 34, 413–435 (2013).

80. C. A. Farthing, D. H. Sweet, Expression and function of organic cation and anion transporters (SLC22 family) in the CNS. Curr. Pharm. Des. 20, 1472–1486 (2014).

81. A. Nowack, J. Yao, K. L. Custer, S. M. Bajjalieh, SV2 regulates neurotransmitter release via multiple mechanisms. Am. J. Physiol. Cell Physiol. 299, C960–7 (2010).

82. K. Buckley, R. B. Kelly, Identification of a transmembrane glycoprotein specific for secretory vesicles of neural and endocrine cells. J. Cell Biol. 100, 1284–1294 (1985).

83. S. M. Bajjalieh, K. Peterson, R. Shinghal, R. H. Scheller, SV2, a brain synaptic vesicle protein homologous to bacterial transporters. Science 257, 1271–1273 (1992).

84. M. Dong, F. Yeh, W. H. Tepp, C. Dean, E. A. Johnson, R. Janz, E. R. Chapman, SV2 is the protein receptor for botulinum neurotoxin A. Science 312, 592–596 (2006).

85. Z. Vuckovic, P. R. Gentry, A. E. Berizzi, K. Hirata, S. Varghese, G. Thompson, E. T. van der Westhuizen, W. A. C. Burger, R. Rahmani, C. Valant, C. J. Langmead, C. W. Lindsley, J. B. Baell, A. B. Tobin, P. M. Sexton, A. Christopoulos, D. M. Thal, Crystal structure of the M5 muscarinic acetylcholine receptor. Proc. Natl. Acad. Sci. U. S. A. 116, 26001–26007 (2019).

86. S. Passaro, G. Corso, J. Wohlwend, M. Reveiz, S. Thaler, V. R. Somnath, N. Getz, T. Portnoi, J. Roy, H. Stark, D. Kwabi-Addo, D. Beaini, T. Jaakkola, R. Barzilay, Boltz-2: Towards Accurate and Efficient Binding Affinity Prediction, bioRxiv (2025). 10.1101/2025.06.14.659707.

87. J. Cao, M. Spielmann, X. Qiu, X. Huang, D. M. Ibrahim, A. J. Hill, F. Zhang, S. Mundlos, L. Christiansen, F. J. Steemers, C. Trapnell, J. Shendure, The single-cell transcriptional landscape of mammalian organogenesis. Nature 566, 496–502 (2019).

88. S. Siebert, J. A. Farrell, J. F. Cazet, Y. Abeykoon, A. S. Primack, C. E. Schnitzler, C. E. Juliano, Stem cell differentiation trajectories in Hydra resolved at single-cell resolution. Science 365, eaav9314 (2019).

89. D. Z. Faltine-Gonzalez, M. J. Layden, Characterization of nAChRs in Nematostella vectensis supports neuronal and non-neuronal roles in the cnidarian-bilaterian common ancestor. EvoDevo 10, 27 (2019).

90. B. J. Liebeskind, H. A. Hofmann, D. M. Hillis, H. H. Zakon, Evolution of animal neural systems. Annu. Rev. Ecol. Evol. Syst. 48, 377–398 (2017).

91. G. Xing, W.-C. Xiong, L. Mei, Rapsyn as a signaling and scaffolding molecule in neuromuscular junction formation and maintenance. Neurosci. Lett. 731, 135013 (2020).

92. X. Fan, T. Sun, W. Crawford, X. Tan, X. Ou, D. A. Terrar, S. N. Ebert, M. Lei, Pnmt-Derived Cardiomyocytes: Anatomical Localization, Function and Future Perspectives. Front. Physiol. 10, 713 (2019).

93. T. Hiroi, S. Imaoka, Y. Funae, Dopamine formation from tyramine by CYP2D6. Biochem. Biophys. Res. Commun. 249, 838–843 (1998).

94. E. Bromek, A. Haduch, K. Gołembiowska, W. A. Daniel, Cytochrome P450 mediates dopamine formation in the brain in vivo. J. Neurochem. 118, 806–815 (2011).

95. J. Borycz, J. A. Borycz, A. Kubów, V. Lloyd, I. A. Meinertzhagen, Drosophila ABC transporter mutants white, brown and scarlet have altered contents and distribution of biogenic amines in the brain. J. Exp. Biol. 211, 3454–3466 (2008).

96. K. Schmidt, M. Ralle, T. Schaffer, S. Jayakanthan, B. Bari, A. Muchenditsi, S. Lutsenko, ATP7A and ATP7B copper transporters have distinct functions in the regulation of neuronal dopamine-β-hydroxylase. J. Biol. Chem. 293, 20085–20098 (2018).

97. V. Gerbasi, S. Lutsenko, E. J. Lewis, A mutation in the ATP7B copper transporter causes reduced dopamine beta-hydroxylase and norepinephrine in mouse adrenal. Neurochem. Res. 28, 867–873 (2003).

98. L. De Petrocellis, D. Melck, T. Bisogno, A. Milone, V. Di Marzo, Finding of the endocannabinoid signalling system in Hydra, a very primitive organism: possible role in the feeding response. Neuroscience 92, 377–387 (1999).

99. M. Roopin, O. Levy, Melatonin distribution reveals clues to its biological significance in basal metazoans. PLoS One 7, e52266 (2012).

100. H. Weissbach, W. Lovenberg, S. Udenfriend, Characteristics of mammalian histidine decarboxylating enzymes. Biochim. Biophys. Acta 50, 177–179 (1961).

101. J. Yu, M. C. Vogt, B. W. Fox, C. J. J. Wrobel, D. Fajardo Palomino, B. J. Curtis, B. Zhang, H. H. Le, A. Tauffenberger, O. Hobert, F. C. Schroeder, Parallel pathways for serotonin biosynthesis and metabolism in C. elegans. Nat. Chem. Biol., 1–10 (2022).

102. J. H. Welsh, 5-Hydroxytryptamine in coelenterates. Nature 186, 811–812 (1960).

103. I. Hajj-Ali, M. Anctil, Characterization of a serotonin receptor in the cnidarian Renilla koellikeri: a radiobinding analysis. Neurochem. Int. 31, 83–93 (1997).

104. M. D. Davis, S. Kaufman, Products of the tyrosine-dependent oxidation of tetrahydrobiopterin by rat liver phenylalanine hydroxylase. Arch. Biochem. Biophys. 304, 9–16 (1993).

105. S. C. Daubner, J. Melendez, P. F. Fitzpatrick, Reversing the substrate specificities of phenylalanine and tyrosine hydroxylase: aspartate 425 of tyrosine hydroxylase is essential for L-DOPA formation. Biochemistry 39, 9652–9661 (2000).

106. D. B. Fisher, S. Kaufman, Tetrahydropterin oxidation without hydroxylation catalyzed by rat liver phenylalanine hydroxylase. J. Biol. Chem. 248, 4300–4304 (1973).

107. K. Teigen, J. A. McKinney, J. Haavik, A. Martínez, Selectivity and affinity determinants for ligand binding to the aromatic amino acid hydroxylases. Curr. Med. Chem. 14, 455–467 (2007).

108. W. R. Francis, M. Eitel, S. Vargas, M. Adamski, S. H. D. Haddock, S. Krebs, H. Blum, D. Erpenbeck, G. Wörheide, The genome of the contractile demosponge Tethya wilhelma and the evolution of metazoan neural signalling pathways, bioRxiv (2017). 10.1101/120998.

109. X. Xiang, A. A. Vilar Gomez, S. P. Blomberg, H. Yuan, B. M. Degnan, S. M. Degnan, Potential for host-symbiont communication via neurotransmitters and neuromodulators in an aneural animal, the marine sponge Amphimedon queenslandica. Front. Neural Circuits 17, 1250694 (2023).

110. O. I. Kravchuk, A. D. Finoshin, Y. O. Nikishina, V. I. Melnikova, I. V. Kublanov, D. A. Sutormin, A. N. Rusanova, M. T. Ri, A. B. Isaev, K. V. Mikhailov, R. H. Ziganshin, K. I. Adameyko, A. A. Anashkina, V. M. Ignatyuk, N. G. Gornostaev, E. E. Voronezhskaya, A. M. Sokolova, V. S. Mikhailov, Y. V. Lyupina, Evolutionary conservation of dopamine-mediated cellular plasticity in Arctic sponges (Porifera). Front. Mol. Biosci. 12, 1671771 (2025).

111. N. Kaneda, H. Ichinose, K. Kobayashi, K. Oka, F. Kishi, A. Nakazawa, Y. Kurosawa, K. Fujita, T. Nagatsu, Molecular cloning of cDNA and chromosomal assignment of the gene for human phenylethanolamine N-methyltransferase, the enzyme for epinephrine biosynthesis. J. Biol. Chem. 263, 7672–7677 (1988).

112. G. Kass-Simon, P. Pierobon, Cnidarian chemical neurotransmission, an updated overview. Comp. Biochem. Physiol. A Mol. Integr. Physiol. 146, 9–25 (2007).

113. A. Ronowska, A. Szutowicz, H. Bielarczyk, S. Gul-Hinc, J. Klimaszewska-Łata, A. Dyś, M. Zyśk, A. Jankowska-Kulawy, The Regulatory Effects of Acetyl-CoA Distribution in the Healthy and Diseased Brain. Front. Cell. Neurosci. 12, 169 (2018).

114. H. Koepsell, H. Endou, The SLC22 drug transporter family. Pflugers Arch. 447, 666–676 (2004).

115. S. Berrard, H. Varoqui, R. Cervini, M. Israël, J. Mallet, M. F. Diebler, Coregulation of two embedded gene products, choline acetyltransferase and the vesicular acetylcholine transporter. J. Neurochem. 65, 939–942 (1995).

116. L. E. Eiden, The cholinergic gene locus. J. Neurochem. 70, 2227–2240 (1998).

117. M. Goulty, C. Lewis, F. Nicolini, R. Khalid, H. Syed, P. Oliveri, M. Thomas, E. Rosato, R. Feuda, Monoaminergic neurons share transcriptional identity across Bilaterian animals, bioRxiv (2025). 10.1101/2025.10.10.679534.

118. S. R. Eddy, Accelerated Profile HMM Searches. PLoS Comput. Biol. 7, e1002195 (2011).

119. A. J. Enright, S. Van Dongen, C. A. Ouzounis, An efficient algorithm for large-scale detection of protein families. Nucleic Acids Res. 30, 1575–1584 (2002).

120. K. Katoh, K. Misawa, K.-I. Kuma, T. Miyata, MAFFT: a novel method for rapid multiple sequence alignment based on fast Fourier transform. Nucleic Acids Res. 30, 3059–3066 (2002).

121. L.-T. Nguyen, H. A. Schmidt, A. von Haeseler, B. Q. Minh, IQ-TREE: a fast and effective stochastic algorithm for estimating maximum-likelihood phylogenies. Mol. Biol. Evol. 32, 268–274 (2015).

122. D. T. Hoang, O. Chernomor, A. von Haeseler, B. Q. Minh, L. S. Vinh, UFBoot2: Improving the Ultrafast Bootstrap Approximation. Mol. Biol. Evol. 35, 518–522 (2018).

123. J. Huerta-Cepas, F. Serra, P. Bork, ETE 3: Reconstruction, Analysis, and Visualization of Phylogenomic Data. Mol. Biol. Evol. 33, 1635–1638 (2016).

124. F. Sievers, A. Wilm, D. Dineen, T. J. Gibson, K. Karplus, W. Li, R. Lopez, H. McWilliam, M. Remmert, J. Söding, J. D. Thompson, D. G. Higgins, Fast, scalable generation of high-quality protein multiple sequence alignments using Clustal Omega. Mol. Syst. Biol. 7, 539 (2011).

125. M. N. Price, P. S. Dehal, A. P. Arkin, FastTree 2 – approximately maximum-likelihood trees for large alignments. PLoS One 5, e9490 (2010).

126. H. Li, J. Janssens, M. De Waegeneer, S. S. Kolluru, K. Davie, V. Gardeux, W. Saelens, F. P. A. David, M. Brbić, K. Spanier, J. Leskovec, C. N. McLaughlin, Q. Xie, R. C. Jones, K. Brueckner, J. Shim, S. G. Tattikota, F. Schnorrer, K. Rust, T. G. Nystul, Z. Carvalho-Santos, C. Ribeiro, S. Pal, S. Mahadevaraju, T. M. Przytycka, A. M. Allen, S. F. Goodwin, C. W. Berry, M. T. Fuller, H. White-Cooper, E. L. Matunis, S. DiNardo, A. Galenza, L. E. O’Brien, J. A. T. Dow, FCA Consortium§, H. Jasper, B. Oliver, N. Perrimon, B. Deplancke, S. R. Quake, L. Luo, S. Aerts, D. Agarwal, Y. Ahmed-Braimah, M. Arbeitman, M. M. Ariss, J. Augsburger, K. Ayush, C. C. Baker, T. Banisch, K. Birker, R. Bodmer, B. Bolival, S. E. Brantley, J. A. Brill, N. C. Brown, N. A. Buehner, X. T. Cai, R. Cardoso-Figueiredo, F. Casares, A. Chang, T. R. Clandinin, S. Crasta, C. Desplan, A. M. Detweiler, D. B. Dhakan, E. Donà, S. Engert, S. Floc’hlay, N. George, A. J. González-Segarra, A. K. Groves, S. Gumbin, Y. Guo, D. E. Harris, Y. Heifetz, S. L. Holtz, F. Horns, B. Hudry, R.-J. Hung, Y. N. Jan, J. S. Jaszczak, G. S. X. E. Jefferis, J. Karkanias, T. L. Karr, N. S. Katheder, J. Kezos, A. A. Kim, S. K. Kim, L. Kockel, N. Konstantinides, T. B. Kornberg, H. M. Krause, A. T. Labott, M. Laturney, R. Lehmann, S. Leinwand, J. Li, J. S. S. Li, K. Li, K. Li, L. Li, T. Li, M. Litovchenko, H.-H. Liu, Y. Liu, T.-C. Lu, J. Manning, A. Mase, M. Matera-Vatnick, N. R. Matias, C. E. McDonough-Goldstein, A. McGeever, A. D. McLachlan, P. Moreno-Roman, N. Neff, M. Neville, S. Ngo, T. Nielsen, C. E. O’Brien, D. Osumi-Sutherland, M. N. Özel, I. Papatheodorou, M. Petkovic, C. Pilgrim, A. O. Pisco, C. Reisenman, E. N. Sanders, G. Dos Santos, K. Scott, A. Sherlekar, P. Shiu, D. Sims, R. V. Sit, M. Slaidina, H. E. Smith, G. Sterne, Y.-H. Su, D. Sutton, M. Tamayo, M. Tan, I. Tastekin, C. Treiber, D. Vacek, G. Vogler, S. Waddell, W. Wang, R. I. Wilson, M. F. Wolfner, Y.-C. E. Wong, A. Xie, J. Xu, S. Yamamoto, J. Yan, Z. Yao, K. Yoda, R. Zhu, R. P. Zinzen, Fly Cell Atlas: A single-nucleus transcriptomic atlas of the adult fruit fly. Science 375, eabk2432 (2022).

127. A. Sebé-Pedrós, B. Saudemont, E. Chomsky, F. Plessier, M.-P. Mailhé, J. Renno, Y. Loe-Mie, A. Lifshitz, Z. Mukamel, S. Schmutz, S. Novault, P. R. H. Steinmetz, F. Spitz, A. Tanay, H. Marlow, Cnidarian Cell Type Diversity and Regulation Revealed by Whole-Organism Single-Cell RNA-Seq. Cell 173, 1520–1534.e20 (2018).

128. A. Sebé-Pedrós, E. Chomsky, K. Pang, D. Lara-Astiaso, F. Gaiti, Z. Mukamel, I. Amit, A. Hejnol, B. M. Degnan, A. Tanay, Early metazoan cell type diversity and the evolution of multicellular gene regulation. Nat Ecol Evol 2, 1176–1188 (2018).

129. J. M. Musser, K. J. Schippers, M. Nickel, G. Mizzon, A. B. Kohn, C. Pape, P. Ronchi, N. Papadopoulos, A. J. Tarashansky, J. U. Hammel, F. Wolf, C. Liang, A. Hernández-Plaza, C. P. Cantalapiedra, K. Achim, N. L. Schieber, L. Pan, F. Ruperti, W. R. Francis, S. Vargas, S. Kling, M. Renkert, M. Polikarpov, G. Bourenkov, R. Feuda, I. Gaspar, P. Burkhardt, B. Wang, P. Bork, M. Beck, T. R. Schneider, A. Kreshuk, G. Wörheide, J. Huerta-Cepas, Y. Schwab, L. L. Moroz, D. Arendt, Profiling cellular diversity in sponges informs animal cell type and nervous system evolution. Science 374, 717–723 (2021).

130. S. Levy, A. Elek, X. Grau-Bové, S. Menéndez-Bravo, M. Iglesias, A. Tanay, T. Mass, A. Sebé-Pedrós, A stony coral cell atlas illuminates the molecular and cellular basis of coral symbiosis, calcification, and immunity. Cell 184, 2973–2987.e18 (2021).

131. E. Jurrus, D. Engel, K. Star, K. Monson, J. Brandi, L. E. Felberg, D. H. Brookes, L. Wilson, J. Chen, K. Liles, M. Chun, P. Li, D. W. Gohara, T. Dolinsky, R. Konecny, D. R. Koes, J. E. Nielsen, T. Head-Gordon, W. Geng, R. Krasny, G.-W. Wei, M. J. Holst, J. A. McCammon, N. A. Baker, Improvements to the APBS biomolecular solvation software suite. Protein Sci. 27, 112–128 (2018).

132. M. van Kempen, S. S. Kim, C. Tumescheit, M. Mirdita, C. L. M. Gilchrist, J. Söding, M. Steinegger, Fast and accurate protein structure search with Foldseek. Nat. Biotechnol. 42, 243–246 (2024).

133. A. M. Waterhouse, J. B. Procter, D. M. A. Martin, M. Clamp, G. J. Barton, Jalview Version 2 – a multiple sequence alignment editor and analysis workbench. Bioinformatics 25, 1189–1191 (2009).

134. G. E. Crooks, G. Hon, J.-M. Chandonia, S. E. Brenner, WebLogo: a sequence logo generator. Genome Res. 14, 1188–1190 (2004).

