## Supplemental Data 1 for "From promiscuity to specialization: The origin of cholinergic and monoaminergic signaling is pre-bilaterian"

### Contents

Figures S1–S15 with legends (this file).

Tables S1–S6 are provided as separate Excel files:

**Table S1. Result table for targeted metabolomics.** Values are given in pM, and in ng/mL with and without normalization to sample weight. <LOQ is written when signal was below the limit of quantification. "-" is written when sample was not measured due to insufficient sample volume.

**Table S2. List of species genomic and transcriptomic sequence data used in comparative genomic analyses.** For each species, the name, taxid, number of peptides and data source are provided.

**Table S3. Foldseek search result table of ChAT/CrAT homologs.** The structures of human ChAT (PDB: 2FY3), mouse CrAT (PDB: 2H3P) and ColabFold predictions of multiple non-bilaterian CrAT homologs have been used as input for Foldseek searches (default search parameters, Swissprot as search database). The most similar structure detected, including sequence identity (seq. id), bit score and e-values are shown; hits labeled "AF" are AlphaFold models from the AlphaFold/SwissProt database.

**Table S4. Boltz-2 receptor–ligand binding predictions.** Predicted affinity (log10 IC50, µM) and binder probability (mean of the two affinity heads and per head), together with the structure-confidence metrics (confidence score, pTM, ipTM, ligand ipTM, pLDDT, PDE), for 46 receptors against six ligands; column descriptions are given in the second sheet.

**Table S5. Alignments and phylogenetic trees.** For each tree shown in the figures: protein family, tree (family, clade or subset), number of sequences, alignment method and length, gap filtering, tree-inference method (FastTree with the LG model or IQ-TREE with ModelFinder and ultrafast bootstrap), best-fit model, and the file names in the accompanying data deposit.

**Table S6. Targeted metabolomics acquisition parameters.** Precursor (Q1) and fragment (Q3) masses, expected retention times and compound-dependent parameters (collision energy, cell exit potential, declustering potential) of the scheduled MRM method.

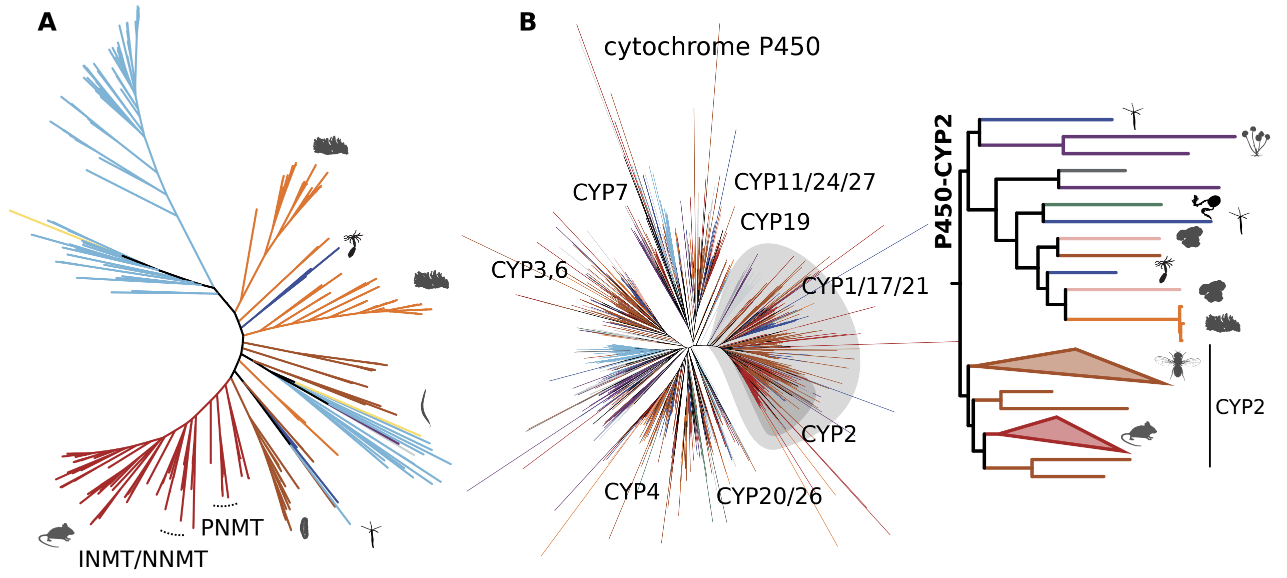


**Figure S1. Phylogenetic trees of the protein families containing the PNMT and cytochrome P450 biosynthetic enzymes.** (**A**) Unrooted ML tree of the NNMT/PNMT/TEMT family. PNMT, INMT, and NNMT have emerged through gene expansion within deuterostomes (dark red). Related proteins are found sparsely in protostome species (brown), cnidarians (blue), sponges (orange), and prokaryotes (bacteria and archaea, cyan and yellow respectively). (**B**) Unrooted ML tree of the cytochrome P450 superfamily (left), and ML tree of the CYP2 family (right) with reduced taxon sampling (see Methods), members of which can catalyze the production of dopamine using tyramine as a substrate. Within CYP2 in Bilateria (bottom part of tree on the right), deuterostome subfamilies have emerged through gene duplications, independently of those in protostomes *D. melanogaster* and *C. elegans* (indicated by the animal silhouettes). Orthologs of the family are found in all animal groups, animal relatives (*Sphaeroforma arctica* in gray) and early diverging fungi (*Allomyces macrogynus* in purple). All color codes are as in Fig. 2.

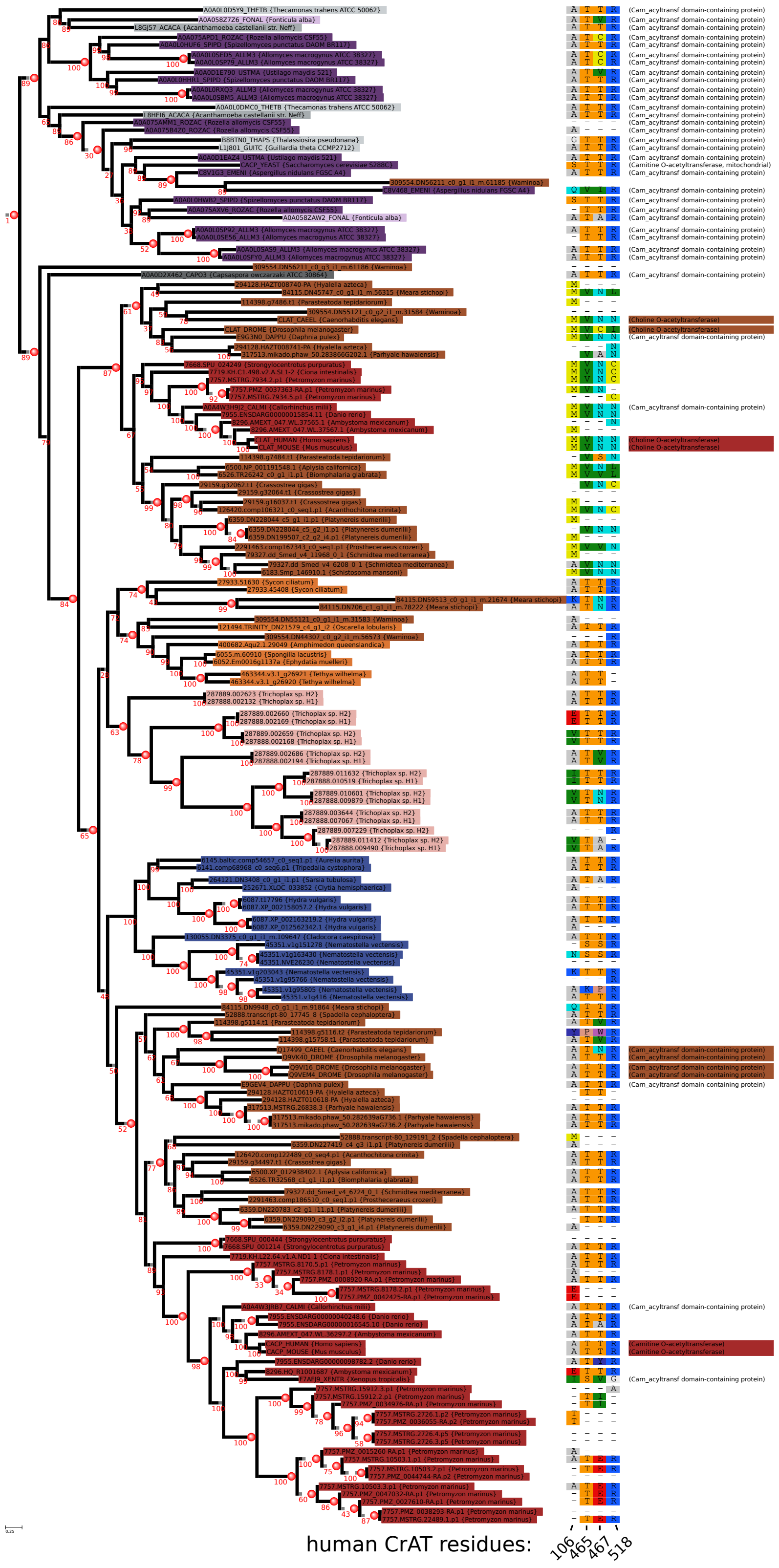


**Figure S2. ML phylogeny of ChAT-CrAT acetyltransferase subfamily and distribution of substrate specificity-defining residues.** ML tree of the ChAT-CrAT clade of Carnitine acetyltransferases with support values and inferred duplication nodes (number in red and red spheres, respectively). The four-residues motif (*51*) is plotted next to the tree tips, and the corresponding positions in the rat and human proteins are shown below each column. Sequence names use the species taxid as a prefix, and the species name is included within “{}”. A strict interpretation of the tree topology would infer that ChAT and CrAT clades derive from a gene duplication in the metazoan common ancestor (ctenophore representatives are missing), and all groups apart from Bilateria lost ChAT. An alternative plausible scenario is that the gene duplication occurred in the bilaterian common ancestor, but cnidarian, placozoan, and sponge sequences are grouped together with CrAT due to high degree of diversification of the ChAT clade.

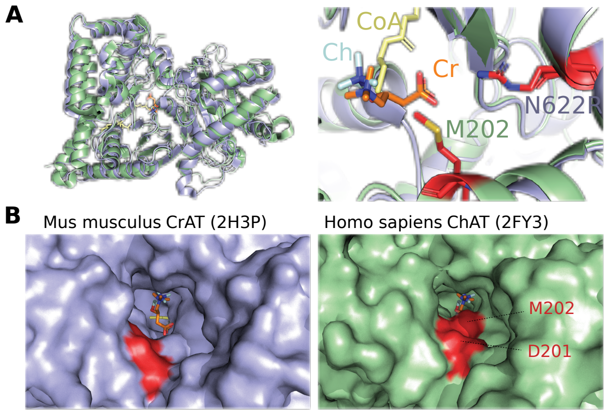


C

| | 201 | 202 | 442 | 554 | 556 | 567 | 569 | 607 | 622 | 629 | 656 | 658 | 668 | 670 | 671 |
|---|---|---|---|---|---|---|---|---|---|---|---|---|---|---|---|
| H.sa.ChAT | D | M | H | Y | S | V | N | Q | N | E | S | S | C | Y | G |
| M.mu.CrAT | T | A | H | Y | S | T | T | H | R | L | S | S | M | F | G |
| S.la.CrAT | A | A | H | Y | S | T | T | H | R | L | S | S | L | F | G |
| H.vu.t17796aep | C | A | H | Y | T | T | T | H | R | M | S | S | L | F | G |
| H.vu.X_002158057.2 | C | A | H | Y | T | T | T | H | R | M | S | S | L | F | G |
| H.vu.X_002163219.2 | Y | A | H | F | A | T | T | H | R | L | C | S | V | F | A |
| N.ve.v1g95805 | V | A | H | K | C | I | N | H | R | L | S | S | V | F | G |
| N.ve.v1g163430 | - | N | H | Y | T | S | S | H | R | C | S | S | T | F | G |
| N.ve.v1g203043 | V | K | H | H | A | T | T | H | R | L | S | R | V | I | R |
| N.ve.v1g416 | I | A | Y | Y | T | T | T | H | R | L | S | S | - | - | - |

charged negative · charged positive · uncharged polar · uncharged hydrophobic

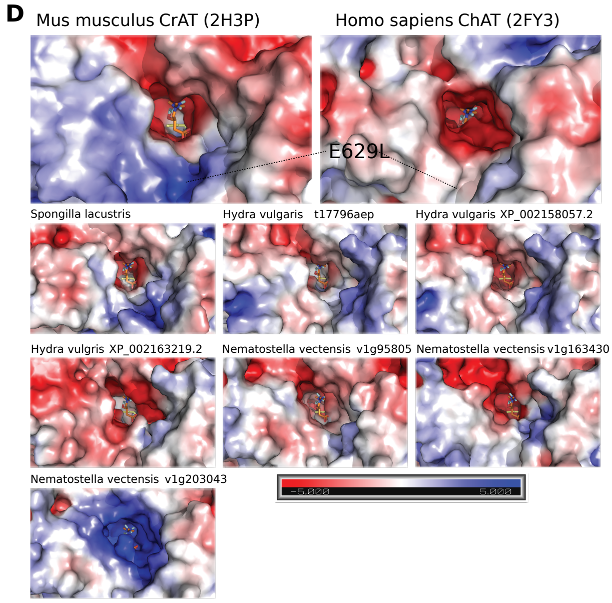


**Figure S3. Structure and sequence comparison between different CrAT and ChAT homologs.** (**A**) The left panel shows the structural alignment of human ChAT (green) (PDB: 2FY3) and mouse CrAT (blue) (PDB: 2H3P) with an RMSD of 1.24 Å. The right panel shows part of the active site of ChAT/CrAT with choline (light blue), carnitine (orange) and CoA (yellow) bound. Red residues are important in distinguishing ChAT and CrAT active sites. Methionine M202, present only in ChATs, sterically prevents the binding of larger substrates such as carnitine. Arginine at position 622 (R518 in mouse CrAT; N622 in ChATs), present only in CrATs, electrically favors the binding of negatively charged carnitine while repelling positively charged choline. (**B**) Comparison of substrate entrance channels between mouse CrAT and human ChAT. Red residues are specifically important in distinguishing CrAT and ChAT substrate channels. Aspartic acid D201, present only in ChATs, repels negatively charged substrates such as carnitine from entering the active site of ChATs, increasing its substrate specificity. (**C**) Comparison of divergent residues between human ChAT and multiple bilaterian and non-bilaterian CrAT homologs. Residues were collected from the literature (48–52) and from Sequence Harmony calculations. Residue numbering is derived from human ChAT. (**D**) Comparison of electrostatic surface potentials of ChAT/CrAT substrate channels. Non-bilaterian structures were predicted using ColabFold. Surface potentials were calculated using the APBS Electrostatics plugin (*131*) of PyMOL (v2.3.5; Schrodinger, Inc.). Red color indicates negative surface potential, blue color indicates positive potential. As expected from the sequence comparison, human ChAT exhibits enhanced negative potential, allowing the binding of positively charged choline. The substitution of leucine by glutamic acid (E629) in ChATs changes the electrostatic potential of the entry channel significantly in favor of positively charged amino acids, excluding negatively charged carnitine. Non-bilaterian CrATs exhibit similar charges to mouse CrAT. v1g203043 features a highly positively charged binding pocket relative to all other enzymes. This excludes positively charged choline from binding, however its substrate specificity remains to be resolved.

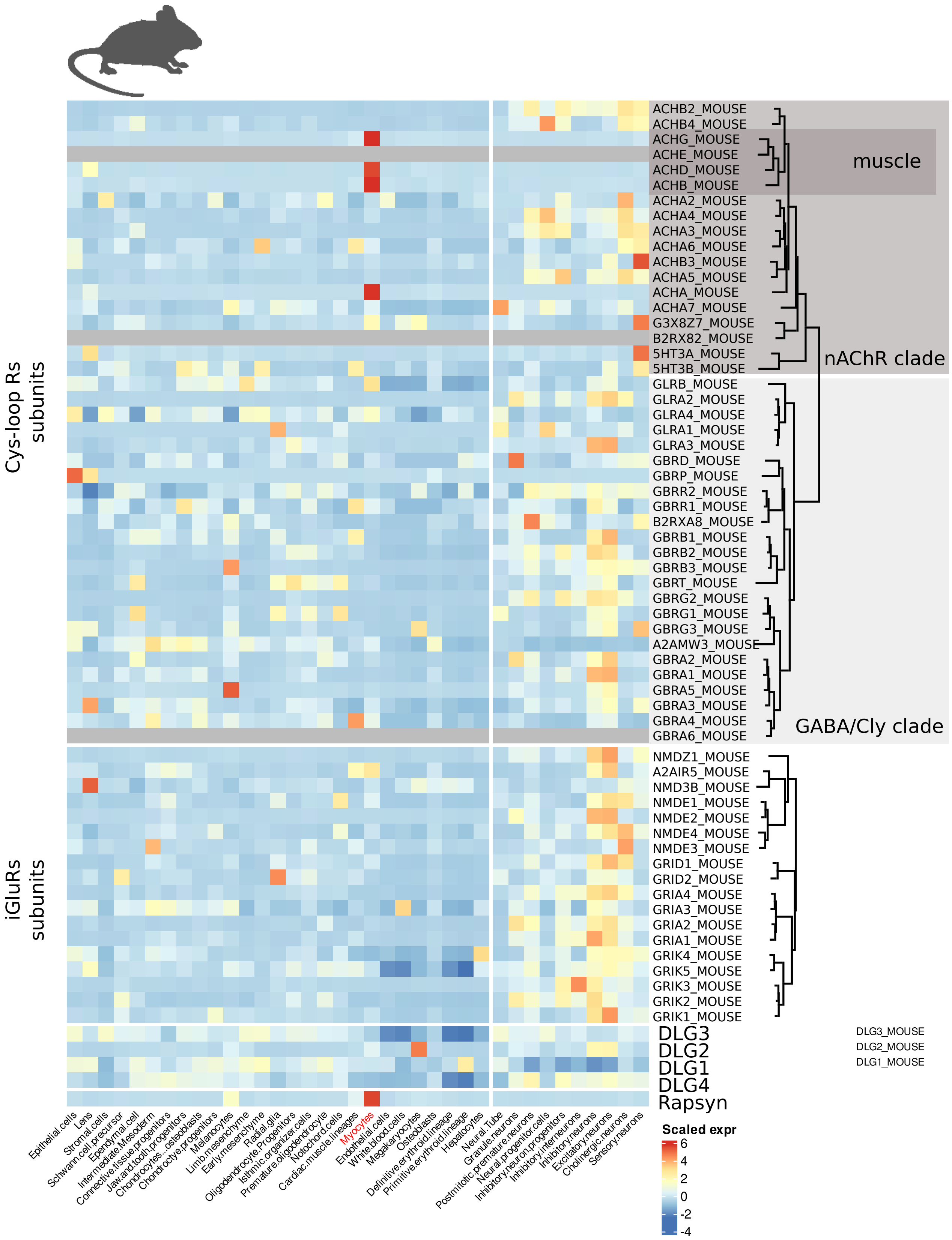


**Figure S4. Single-cell expression of ionotropic Cys-loop and glutamate receptors channel subunits in *Mus musculus.*** Heatmap of the normalized expression values of the genes within each family. The trees on the right derive from ML phylogenies and indicate the relationships between the subunits within each family.

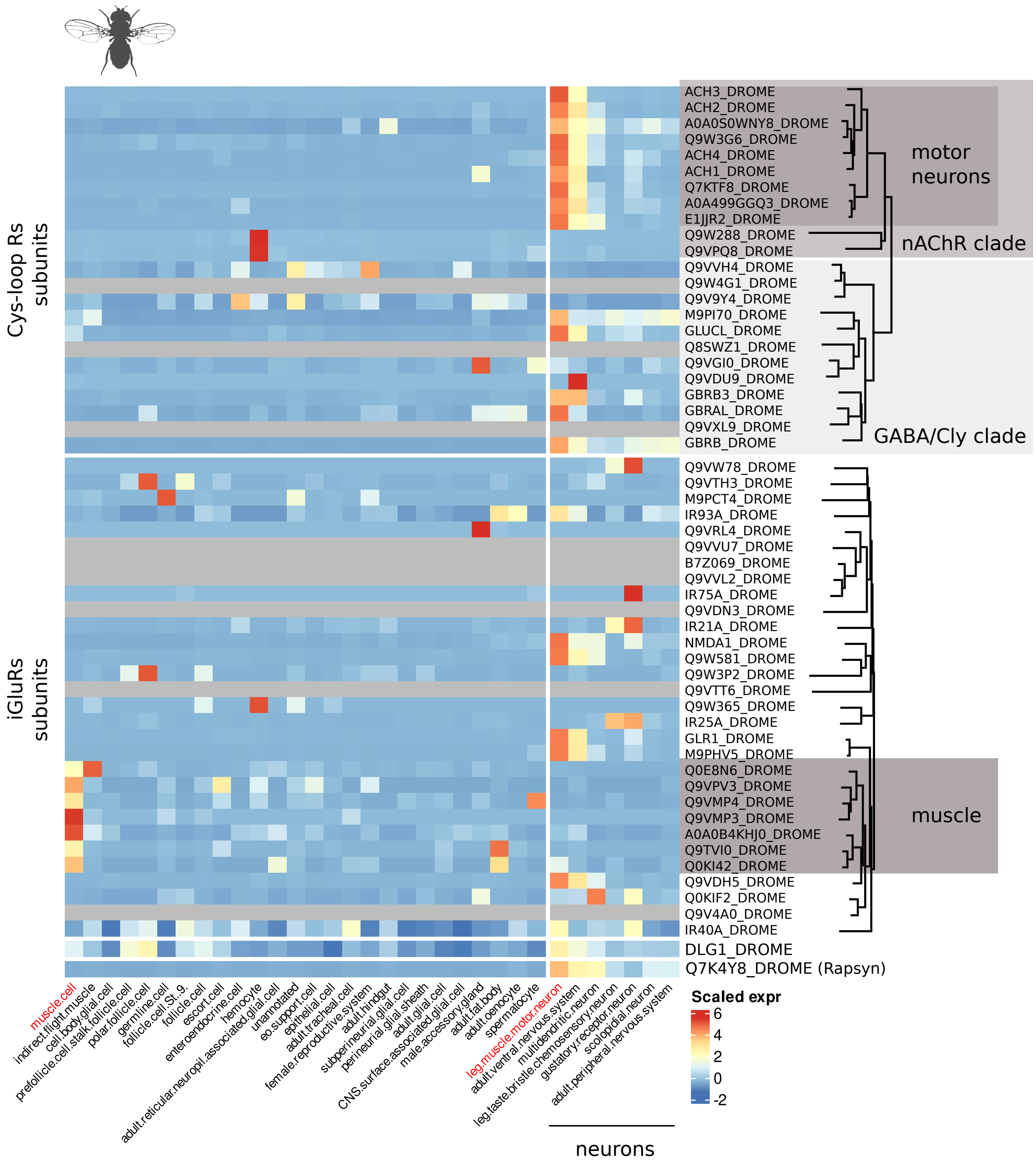


**Figure S5. Single-cell expression of ionotropic Cys-loop and glutamate receptors channel subunits in *Drosophila melanogaster.***

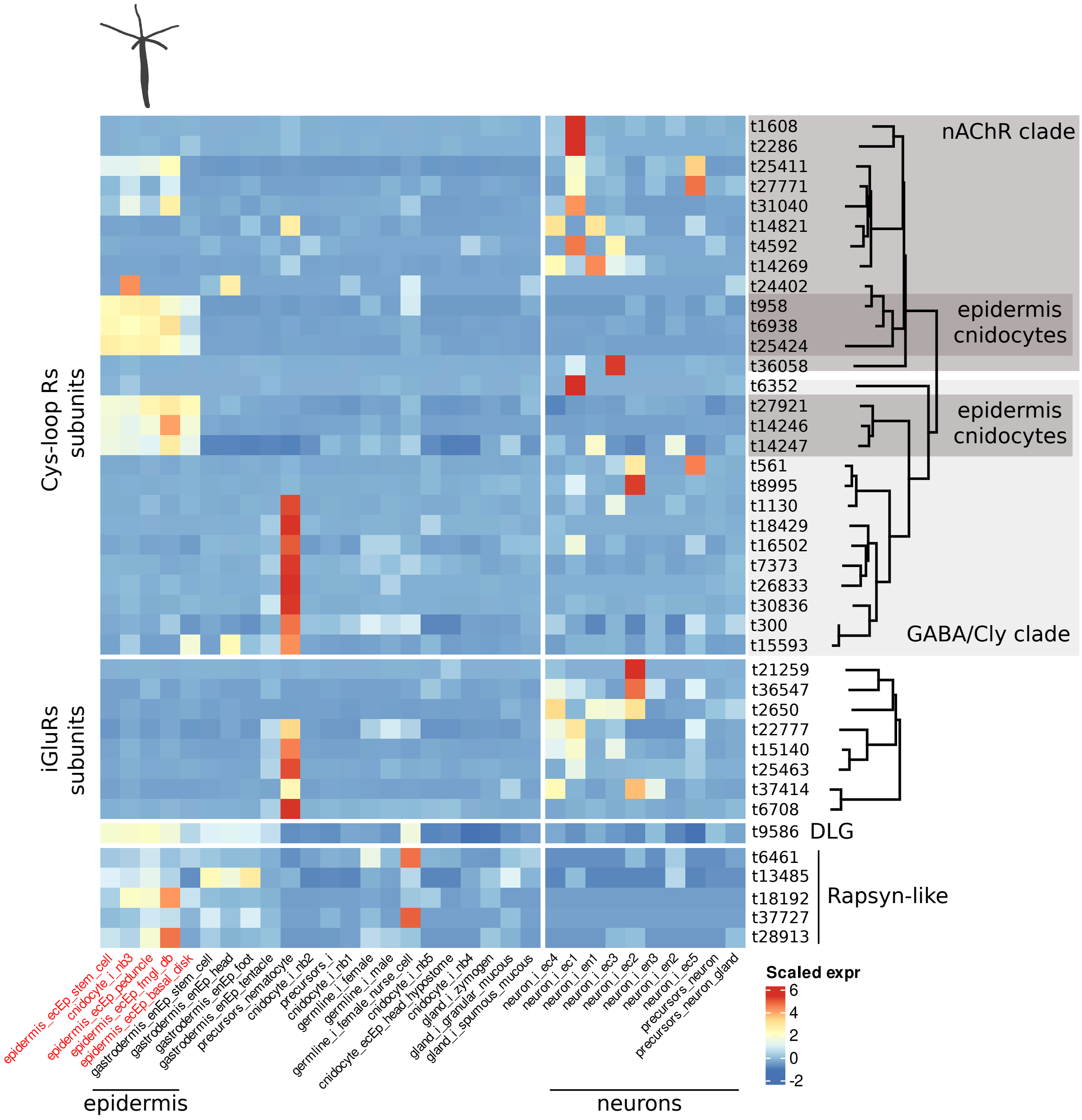


**Figure S6. Single-cell expression of ionotropic Cys-loop and glutamate receptors channel subunits in *Hydra vulgaris*.**

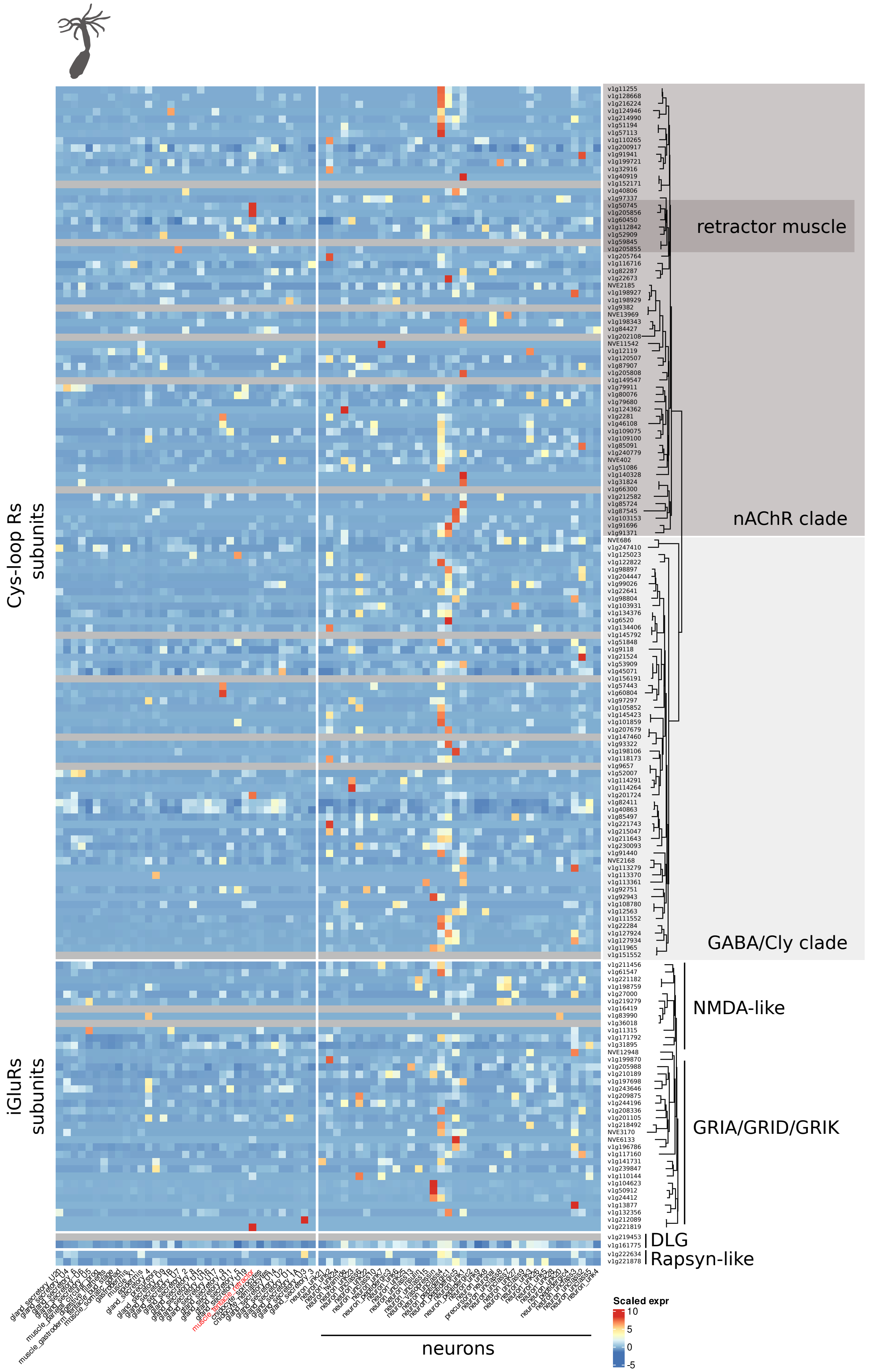


**Figure S7. Single-cell expression of ionotropic Cys-loop and glutamate receptors channel subunits in *Nematostella vectensis.***

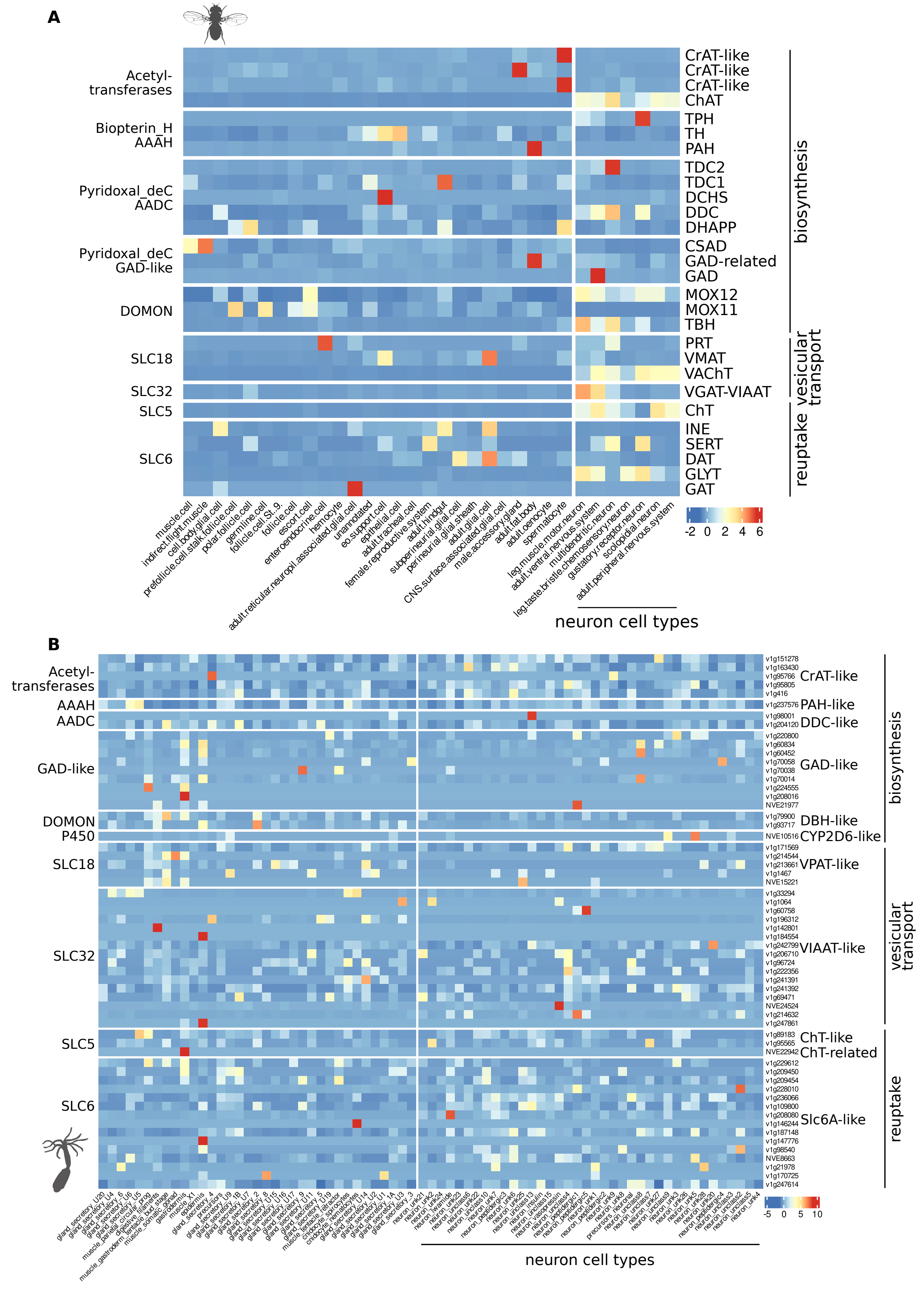


**Figure S8. Single-cell expression of biosynthesis enzymes, vesicular, and reuptake transporter genes in (A) *D. melanogaster* and (B) *N. vectensis*.** All details are as in Fig. 7.

**A**

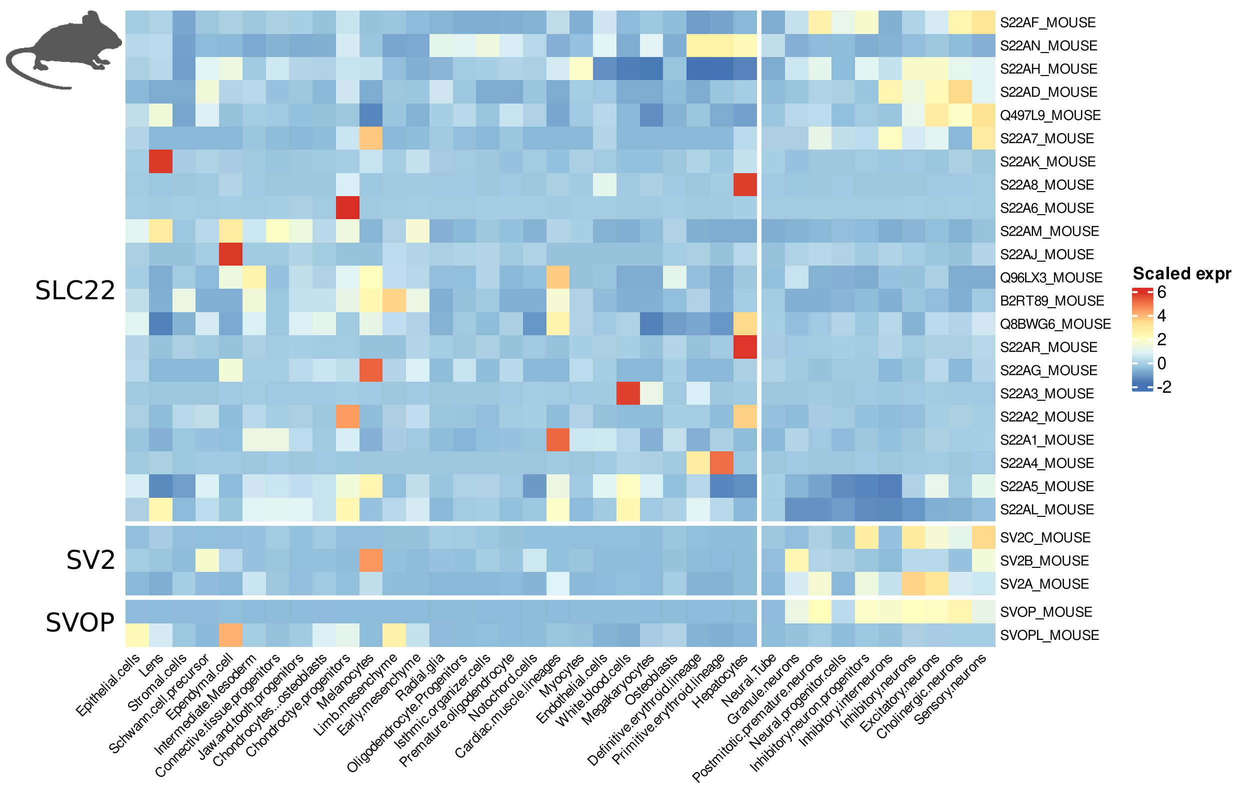


**B**

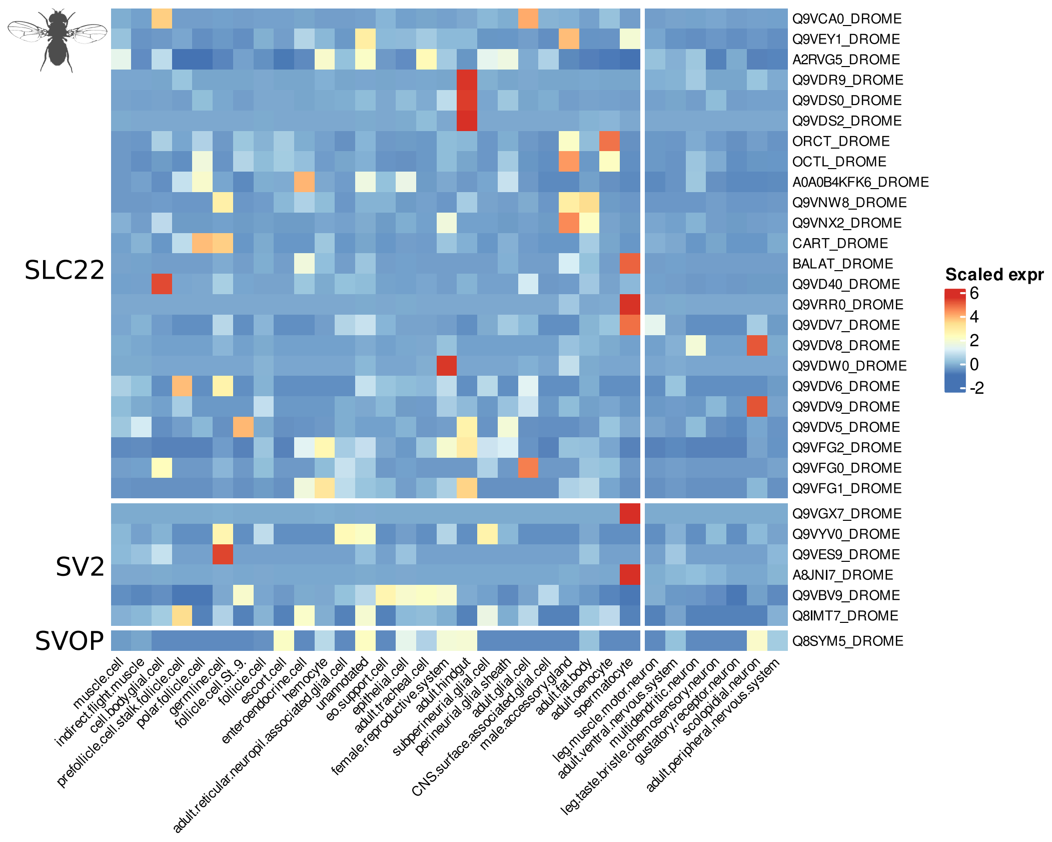


**Figure S9. Single-cell expression of the SLC22 MFS transporters in the bilaterians (A) *M. musculus* and** (B) *D. melanogaster*.

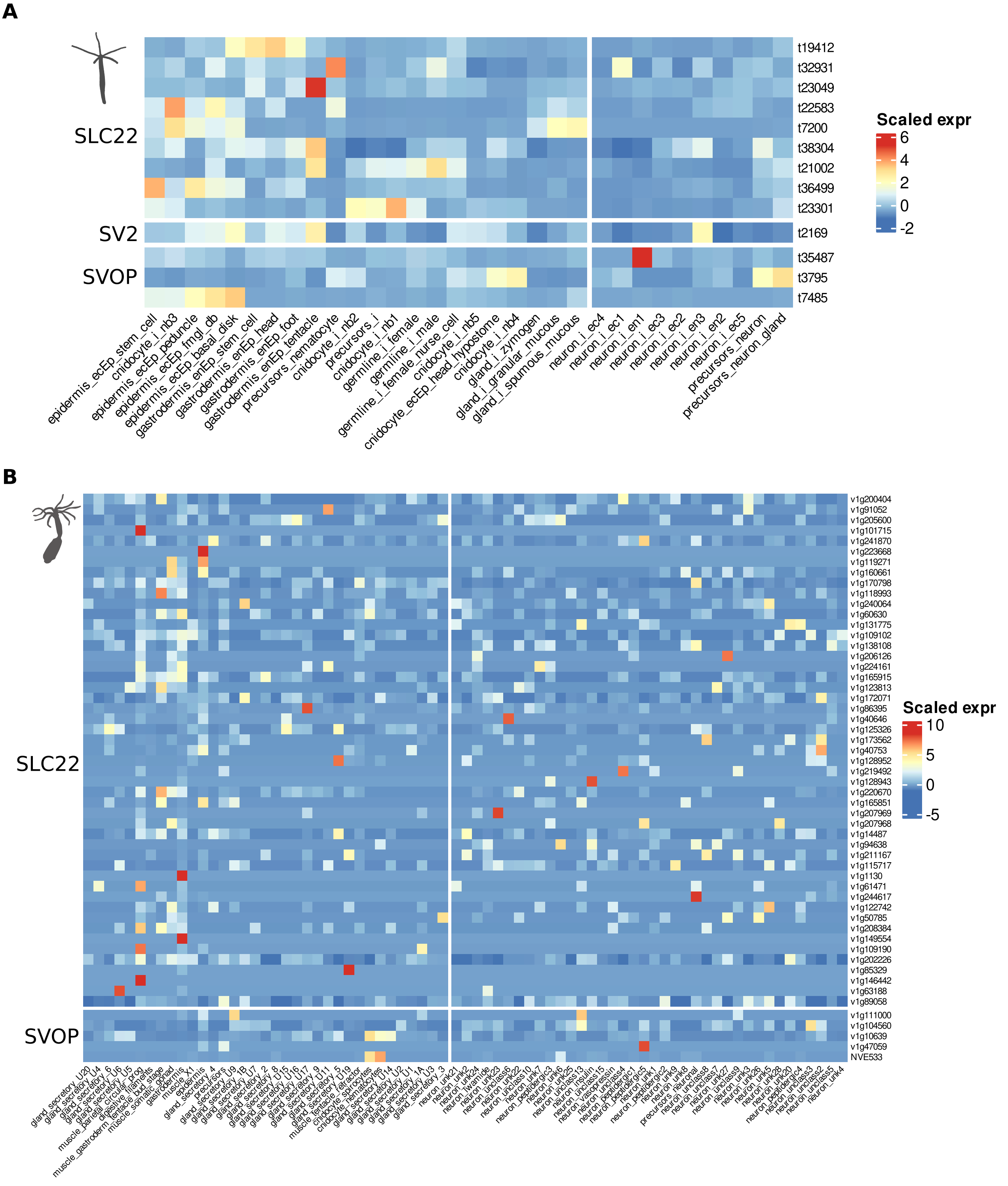


**Figure S10. Single-cell expression of the SLC22 MFS transporters in the cnidarians (A)** ***H.*** *vulgaris* and (B) *N. vectensis***.**

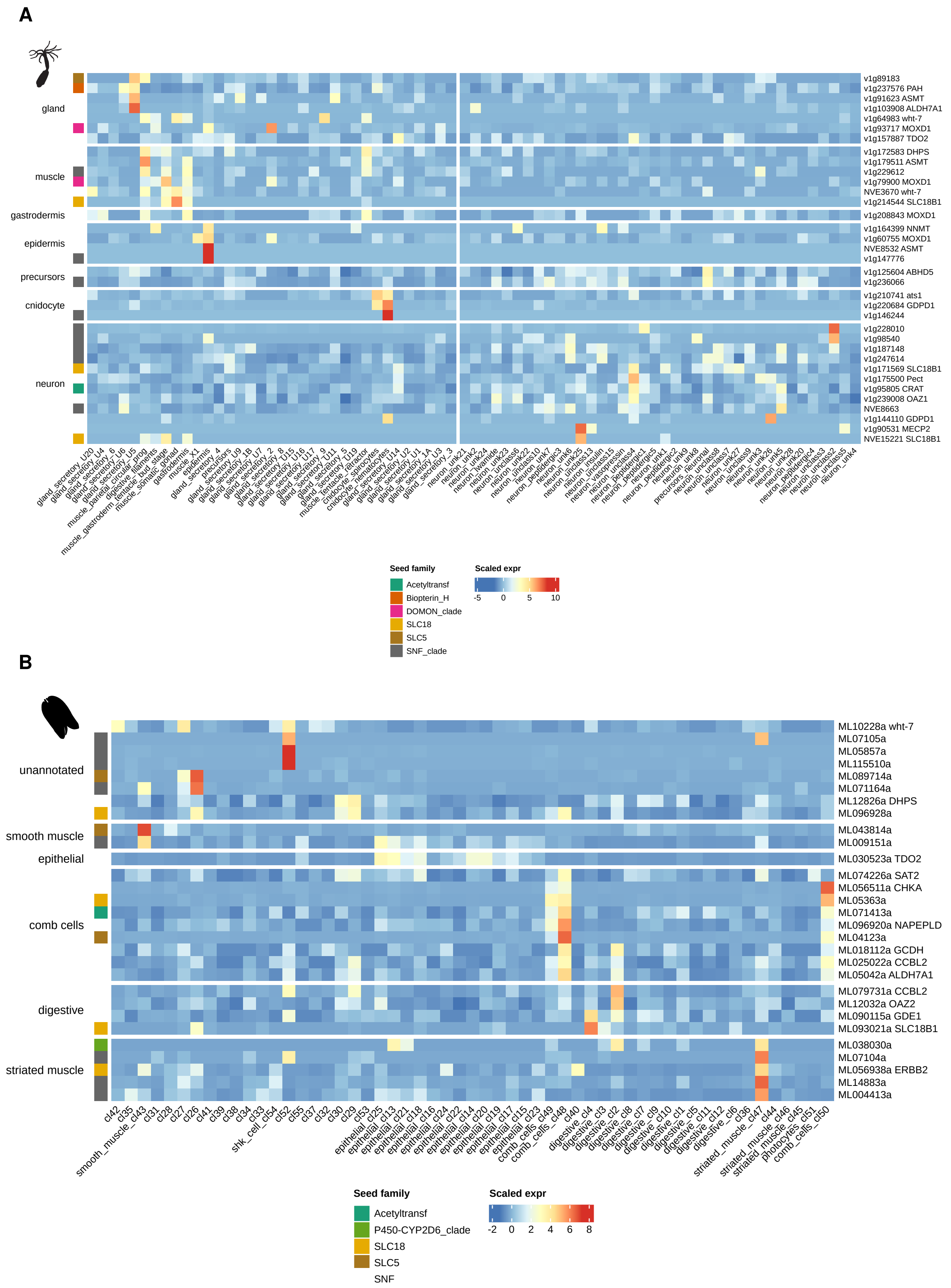


**Figure S11. Monoamine metabolism-associated coregulons in the neuron-bearing non-bilaterians *Nematostella vectensis* and *Mnemiopsis leidyi*.** (A) *N. vectensis*. (B) *M. leidyi*. Genes are grouped by the cell-type class of the cell type with peak expression; all other details are as in Fig. 8.

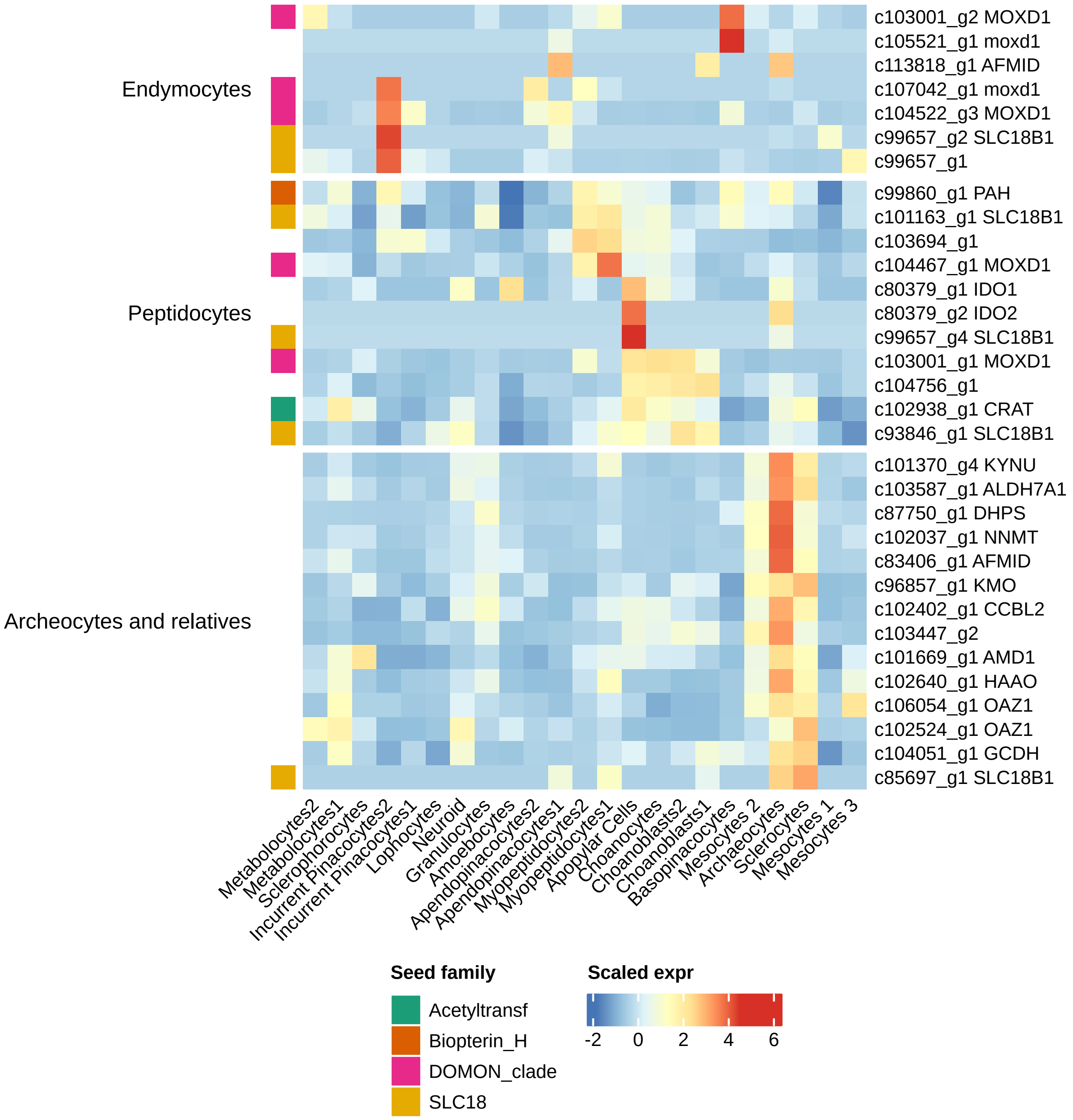


**Figure S12. Monoamine metabolism-associated coregulon in *Spongilla lacustris*.** Genes are grouped by the cell-type family of the cell type with peak expression; all other details are as in Fig. 8.

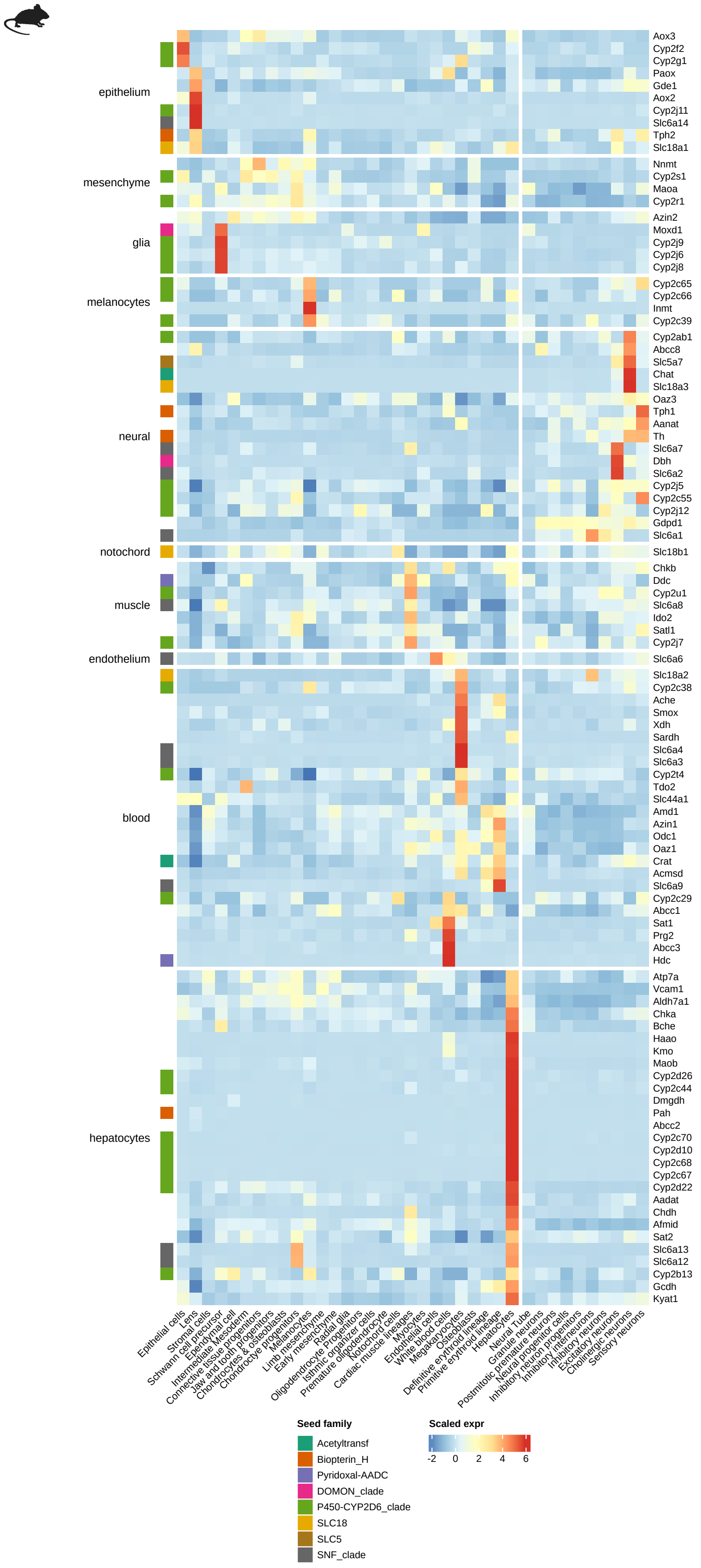


**Figure S13. Monoamine metabolism-associated coregulon in *Mus musculus*.** Genes are grouped by the lineage of the cell type with peak expression; all other details are as in Fig. 8. The coregulon is dominated by lineage-specific CYP2 P450 expansions (CYP2D6, CYP2J2, CYP2C8, CYP2C18, CYP2F1, CYP2B6 and several paralogs of each), together with the dedicated monoamine machinery TPH1, SLC6A2/NET and SLC18A1/VMAT1.

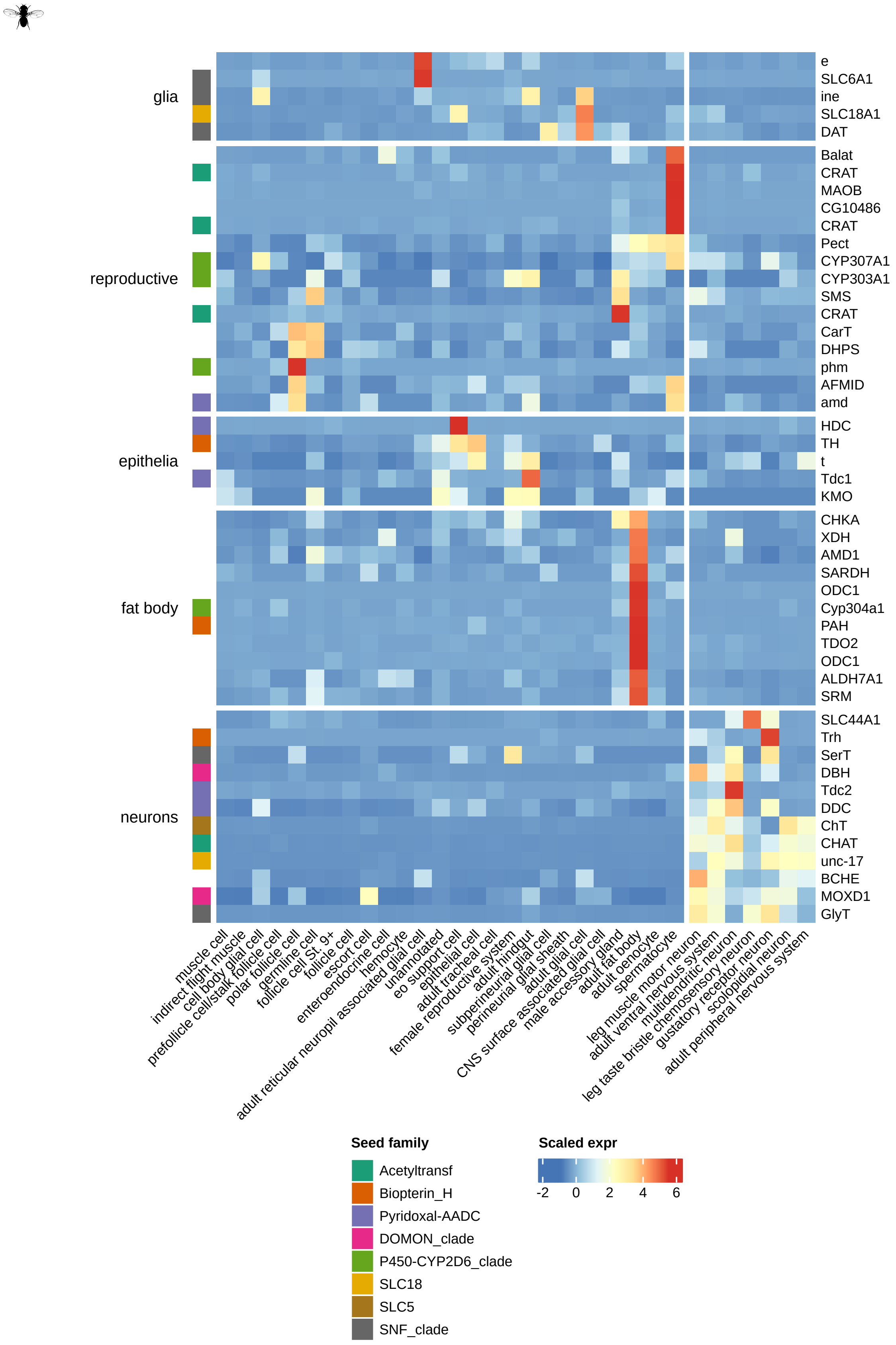


**Figure S14. Monoamine metabolism-associated coregulon in *Drosophila melanogaster*.** Genes are grouped by the tissue of the cell type with peak expression; all other details are as in Fig. 8. The coregulon contains the dedicated insect specialists Tβh (octopamine), Tdc2 (tyramine/octopamine), Trh (serotonin) and Ddc, the cholinergic effectors ChAT, VAChT, ChT and Ace, the dopamine transporter DAT and the vesicular monoamine transporter Vmat, a complete polyamine-biosynthesis pathway (SamDC, Odc1, Odc2, SpdS, Sms) and three kynurenine-pathway enzymes (vermilion/TDO2, cinnabar/KMO, kynurenine formamidase/AFMID).

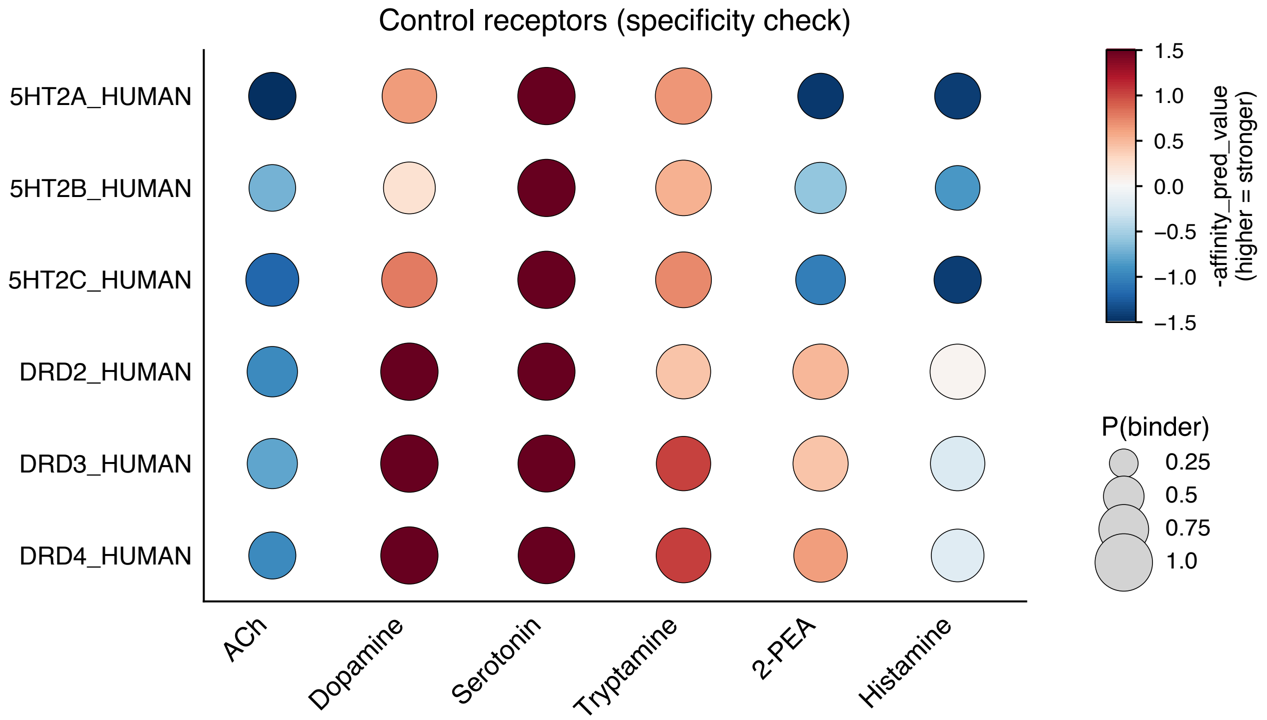


**Figure S15. Boltz-2 binding-affinity predictions for bilaterian serotonin and dopamine receptors.** Human 5-HT2A–C and DRD2–4 receptors against the six candidate ligands; dot size indicates the predicted binding probability and color the predicted affinity, as in Fig. 4B. All predictions are listed in Table S4.
